# H3K4me1-marked Enhancer Activation in Resistant Prostate Cancers Implicates SOX4 and MENIN Inhibition as Therapeutic Strategies

**DOI:** 10.1101/2021.09.02.458625

**Authors:** Christopher J. Terranova, Mayinuer Maitituoheti, Ayush T. Raman, Ming Tang, Nazanin Esmaeili Anvar, Archit K. Ghosh, Samir B. Amin, Jonathan Schulz, Elias Orouji, Filippo Giancotti, Patricia Troncoso, Christopher J. Logothetis, Timothy Thompson, Kunal Rai

**Author notes:** These authors contributed equally (Christopher J. Terranova, Mayinuer Maitituoheti).

## Abstract

Chromatin elements and regulators play important roles during progression of prostate cancer, however, their involvement in response to therapy is less well understood. Using comprehensive chromatin profiling of patient-derived tumors, we find that enhancer elements marked by H3K4me1 are highly enriched in aggressive therapy-resistant prostate cancers on important resistance-driving genes, such as those involved in FOXA1, NOTCH and TGF-β signaling. Importantly, by targeting H3K4me1-elements through inhibition of the MLL complex, a H3K4 methyltransferase, we reduced the proliferative capacity and H3K4me1-associated loci in enzalutamide-resistant prostate cancer lines. We identify AR, FOXA1, HOXB13 and SOX4 as a subset of core TFs that are critical for establishing transcriptional networks via active enhancer reprogramming during acquisition of resistance to therapy. Knock-down of SOX4 reduced cell proliferation and disrupted the H3K4me1 enhancer landscape, further suggesting a role for this TF in therapy-resistance. Overall, our data implicate H3K4me1-marked enhancers as a key epigenetic feature of therapy-resistance, implicate SOX4 in enhancer reprogramming and suggest use of MLL/MENIN inhibitors as a potential therapeutic strategy in high-grade and locally advanced prostate cancers that do not respond to traditional therapies.

## INTRODUCTION

Prostate cancer (PC) is a leading cause of cancer deaths among men in western countries (Siegel et al., 2014). The majority of these deaths occur when localized PC progresses to metastatic castration-resistant prostate cancer (CRPC). Prostate cancer patients who have localized disease often receive treatments through radical prostatectomy or radiotherapy, however, approximately 20%-40% of these cases experience biochemical recurrence (Freedland et al., 2005; Kupelian et al., 2006). The main treatment for recurrence, locally advanced PC, or metastatic spread is androgen deprivation therapy (ADT). ADT consists of surgical or medical castration methods to reduce circulating androgens, however, over time patients eventually develop resistance to ADT and progress to CRPC. FDA approved drugs, such as abiraterone and enzalutamide effectively block androgen synthesis and prevent activation of androgen receptor (AR) respectively (Ryan et al., 2013; Scher et al., 2012). Although these AR targeting agents extend the life of CRPC patients, resistance to treatment is highly recurrent and there is currently no cure for CRPC.

Multiple lines of evidence suggest that aberrant AR signaling is necessary for various aspects of PC development, progression and metastasis (Mohler, 2008; Scher and Sawyers, 2005). Genetic alterations (Waltering et al., 2012),(Brooke and Bevan, 2009)^-^(Chen et al., 2004), oncogenic growth factor signaling (Mulholland et al., 2012) and altered co-regulators (Culig and Santer, 2012) can induce the improper activation of AR even with low amounts of androgens present (Chang et al., 2013; Ishizaki et al., 2013). Recent studies have demonstrated that aberrant enhancer activation of the *AR* gene may play a role in the progression of CRPC (Takeda et al., 2018). Enhancers are long-range regulatory elements that activate transcription by delivering important accessory factors to the promoter region (Calo and Wysocka, 2013). A primary role of enhancer elements is their ability to function as transcription factor (TF) binding platforms (Calo and Wysocka, 2013). During development, enhancer activation generally requires the presence of multiple TFs in order to ensure the activation of lineage-specific genes in response to intrinsic and extrinsic environmental cues. Takeda et al identified a somatically acquired, functionally relevant, intergenic enhancer as a driver of *AR* expression in CRPC (Takeda et al., 2018). Interestingly, the occupation of master TFs on this element, including FOXA1 and HOXB13 (Takeda et al., 2018), suggests this may be a developmental enhancer that becomes reactivated during the selective pressure exerted by antiandrogen treatment.

Similar to AR, the FOXA1 protein also plays a critical role during PC progression and metastatic spread (Parolia et al., 2019; Yang and Yu, 2015). FOXA1/HNF-3α is a pioneer TF that can open compacted chromatin to facilitate the binding of other TFs in promoter and enhancer regions (Gao et al., 2005; Gao et al., 2003; Kaestner, 2010; Kron et al., 2017; Lee et al., 2008). In a subset of TMPRSS2-ERG associated samples, the overexpression of ERG co-opts FOXA1 and HOXB13 to promote an active chromatin landscape associated with increased migration and invasion. Furthermore, genomic rearrangements within the *FOXA1* locus generates a *de novo* enhancer, denoted FOXA1 mastermind (FOXMIND) (Parolia et al., 2019), which drives its own overexpression and that of other oncogenes indicating its central role in mediating PC development and progression. Together, these results suggest PC is governed by a misregulation of enhancer activation and the aberrant binding patterns of core developmental TF such as AR, FOXA1 and HOXB13.

Using combinatorial chromatin state profiling in patient-derived tumors, our study suggests that gain of H3K4me1-marked enhancers is a key epigenetic feature of therapy-resistance in prostate cancer. Targeting these H3K4me1-enhancers through small molecule inhibition of MENIN, a subunit of the Mixed-Lineage Leukemia (MLL) complex (Hughes et al., 2004), disrupted the enhancer and gene expression levels of canonical TF genes, such as *AR,* as well as other developmental TF genes, including *SOX4*. The knock-down of SOX4 decreases cell growth and disrupts the H3K4me1-associated enhancer landscape, further suggesting a functional role for SOX4 in the TF network causing therapy-resistance. Importantly, the ability of MENIN inhibition to disrupt functional H3K4me1-associated enhancers regulating genes known to drive therapy-resistance, such as AR, identifies a plausible mechanism for therapeutically targeting advanced therapy-resistant tumors by blocking H3K4me1-marked elements that cannot be targeted by traditional therapies.

## RESULTS

### The chromatin landscape of therapy-resistant castration resistant prostate cancer

Chromatin state profiling remains a powerful tool for determining the regulatory status of annotated genes and identifying novel elements in non-coding genomic regions. Using ChIP-sequencing for enhancers (H3K27ac and H3K4me1), promoters (H3K4me3), active transcription (H3K79me2), polycomb repression (H3K27me3) and heterochromatin (H3K9me3) repression we describe the cis-regulatory landscape in therapy-resistant prostate cancers. Our cohort was comprised of 4 treatment-naive high-grade locally advanced prostate tumors (radical prostatectomy only; hereby referred as “naive”; and 4 high-grade locally advanced prostate treatment-resistant prostate tumors (hormone ablation therapy and chemotherapy or radiation therapy; hereby referred as “resistant”) (**Figure 1A**). Since chromatin modification based regulation of gene expression is governed by combinatorial patterns of histone modifications (Mikkelsen et al., 2007), we computed multiple chromatin state models (8-states through 30-states) using the ChromHMM algorithm (Ernst and Kellis, 2012) (**Figure 1B** and **S1A-F**). We chose a 15-state model for in-depth analysis as it provided sufficient biological resolution between different chromatin states without any redundancy in states (**Figure S1A-F**). In accordance with the ENCODE Roadmap project (Roadmap Epigenomics et al., 2015), our analysis revealed canonical chromatin state patterns representing both active and repressive domains. These states included active and transcribed promoters within the TSS or flanking regions (E11 and E12); genic and active enhancers within (E13) or outside (E8-E9) the TSS flanking regions; and heterochromatic or polycomb based repression, harboring high levels of either H3K9me3 (E1) or H3K27me3 (E4) respectively (**Figure 1B** and **S2A, B**). Unsupervised clustering analysis of each chromatin state revealed state E8 (active enhancer-H3K4me1 high) and state E10 (weak H3K27ac enhancer) were able to distinguish 3/4 resistant tumors from naive tumors (**Figure S3A-J**). Differential analysis further revealed chromatin state E8 was enriched specifically in resistant tumors (**Figure 1C**) whereas chromatin state E10 displayed enrichment in both naïve and resistant samples (**Figure 1D**). Taken together, these results suggest alterations in H3K4me1-associated active enhancer patterns are the most significant chromatin state difference that occurs in therapy-resistant tumor patient samples in comparison to those tumors that did not need any therapy post-surgery.

**Figure 1:**
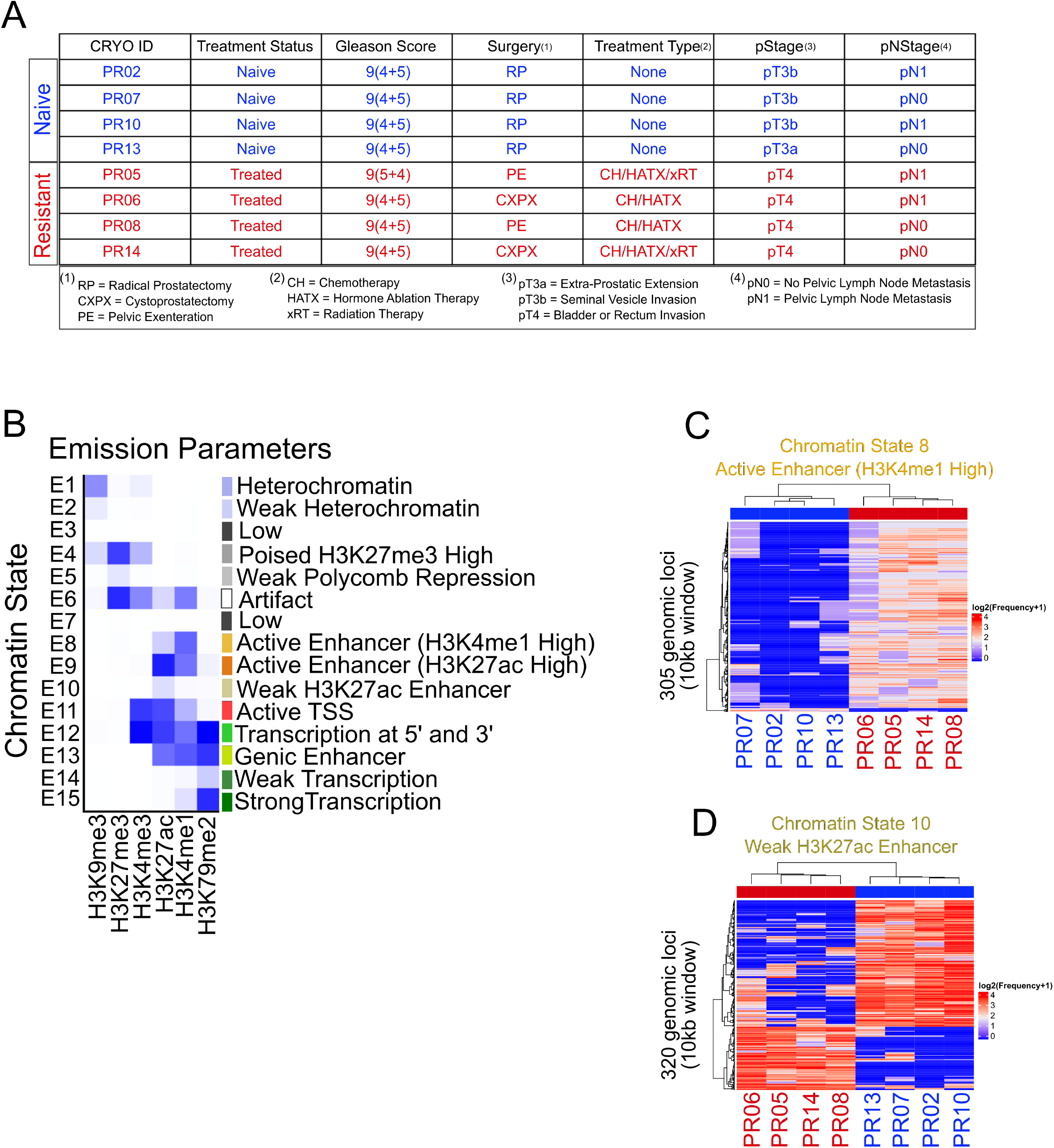
The chromatin landscape of therapy-resistant prostate cancer. A) Pathology overview of naive and resistant prostate tumor samples profiled in this study. B) ChromHMM combinatorial chromatin state patterns identified in 4 naive and 4 resistant prostate tumors based on H3K4me1, H3K27ac, H3K4me3, H3K79me3, H3K27me3 and H3K9me3 histone modifications. C-D) Differential analysis (FDR < 0.05) of chromatin state E8 and D) chromatin state E10 between naive and resistant tumors. Numbers represent total genomic regions based on 10kb bins.

### H3K4me1-associated enhancers are enriched in the therapy-resistant tumors

Since chromatin state E8 contained both active enhancer marks (H3K4me1 and H3K27ac) we next determined whether either of these modifications could distinguish naive and resistant tumors. Differential binding analysis revealed resistant tumors were highly enriched for H3K4me1 compared to that of H3K27ac, with a total 1190 differential sites (p-value ≤ 0.005) compared to 342 sites (p-value ≤ 0.005) respectively (**Figure 2A, B** and **S4A, B**). Consistently, average density analysis and identification of treatment-specific sites demonstrated H3K4me1- and H3K4me1+H3K27ac-associated loci were enriched in both promoter (within −/+5kb of the Transcription Start Site (TSS)) and enhancer (outside −/+5kbTSS) regions in resistant tumors (**Figure 2C-G**, **S4C-F**). Interestingly, this increase was not observed for H3K27ac-associated loci which displayed similar profiles in naive and resistant tumors (**Figure 2C-G**, **S4C-F**). To elucidate the functional role of these gained enhancers, we used Genomic Regions Enrichment of Analysis Tool (GREAT) focusing on active enhancers containing both H3K4me1 and H3K27ac. In naive tumors, active enhancer gains were enriched for pathways including “focal adhesion” and “IL1-mediated signaling events” (**Figure 2H**). In contrast, active enhancer gains in resistant samples were enriched for critical prostate cancer pathways such as “FOXA1 transcription factor network”, “Genes involved in signaling by NOTCH1” and “TGF-beta receptor signaling” (**Figure 2I**). Within these pathways included key regulatory prostate genes such as *AR, FOXA1, HOXB13, CTNNB1* and *MYC*, which displayed gains of H3K4me1-associated enhancers in resistant tumors (**Figure 2J** and **S5A-D**). Together, these results suggest that H3K4me1 enhancers are gained in therapy-resistant prostate tumors and may regulate key genes involved in prostate cancer progression.

**Figure 2:**
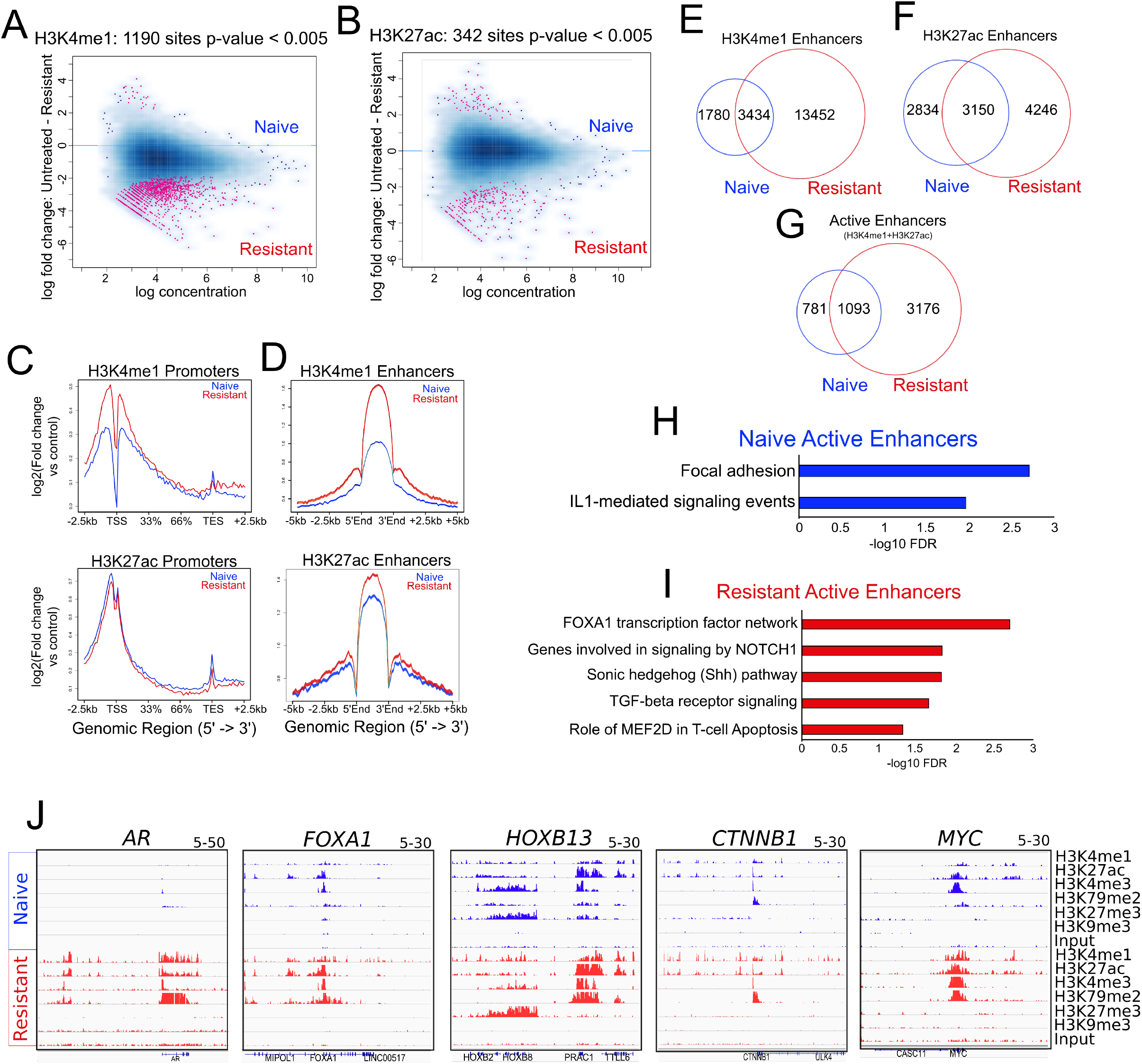
H3K4me1 associated enhancers are enriched in therapy-resistant tumors. A-B) MA plot displaying differential binding sites (log fold change, p-value < 0.005) for H3K4me1 and B) H3K27ac between naive and resistant tumors. C) Average density for H3K4me1 and H3K27ac based on all ensemble promoters (within −/+2.5kbTSS) in naive and resistant samples. D) Average density for H3K4me1 and H3K27ac based on all enhancers (outside −/+5kbTSS) in naive and resistant samples. E-G) Venn diagram analysis of H3K4me1 F) H3K27ac and G) active enhancer (H3K4me1+H3K27ac) peaks in naive and resistant tumors. H-I) Top 5 significant GREAT MSigDB Pathways based on active enhancer peaks (H3K4me1+H3K27ac) in naive tumors and I) resistant tumors. Hyper FDR q-values are displayed as −log10. J) ChIP-seq supertracks displaying H3K4me1, H3K27ac, H3K4me3, H3K79me2, H3K27me3 and H3K9me3 histone modifications on the *AR, FOXA1, HOXB13, CTNNB1* and *MYC* loci in naive (blue) and resistant (red) tumors.

### Inhibition of MLL complex reduces H3K4me1-associated enhancers interacting with critical prostate cancer genes

To determine the functional consequence of H3K4me1-associated enhancer gains, we next focused on potential therapeutic implications of our findings. Previous studies have demonstrated small molecules targeting MLL-MENIN binding surface, such as MI-503, blocks AR signaling and the growth of castration resistant tumors in mice (Malik et al., 2015). As the MLL complex is a well-known histone methyltransferase for H3K4me1 (Calo and Wysocka, 2013), we next determined whether MI-503, a MENIN inhibitor (Malik et al., 2015), can alter the binding pattern of H3K4me1 in enzalutamide-resistant cell line 22RV1. Treatment with MI-503, but not enzalutamide (ENZ) alone, decreased cell proliferation in 22RV1 cells, that serve as a model of ENZ resistance **(Figure 3A and Figure S6A).** ChIP-seq analysis further revealed that MI-503 reduced the number of top variable regions of H3K4me1-associated loci, which was not observed for H3K27ac-associated loci (**Figure 3B, C**). Consistently, MI-503 treatment decreased the average density of H3K4me1 in both promoter and enhancer regions (**Figure 3D, E**). To determine how MLL inhibition impacts gene expression we focused specifically on H3K4me1 enhancers that were gained in resistant samples (resistant tumors and 22RV1 cells) and lost upon MLL inhibition (1381 enhancers) (**Figure 3F**). GREAT analysis revealed H3K4me1 enhancer losses were associated with various developmental TFs involved in “Nuclear Receptor transcription”, and “SOX gene families” (Adamo and Ladomery, 2016; Heinlein and Chang, 2004; Kron et al., 2017; Parolia et al., 2019; Takeda et al., 2018; Yang and Yu, 2015) (**Figure 3G** and **Table S1**). Notably, these genes included canonical TFs known to drive prostate carcinogenesis, such as *AR* and *ESRRG*, and those implicated in embryonic development, such as *NR4A2*, *RARB*, *SOX2* and *SOX4* (**Table S1**). Together these results suggest that growth of therapy resistant cells could be abrogated by MENIN inhibitors through the loss of H3K4me1-marked enhancers.

**Figure 3:**
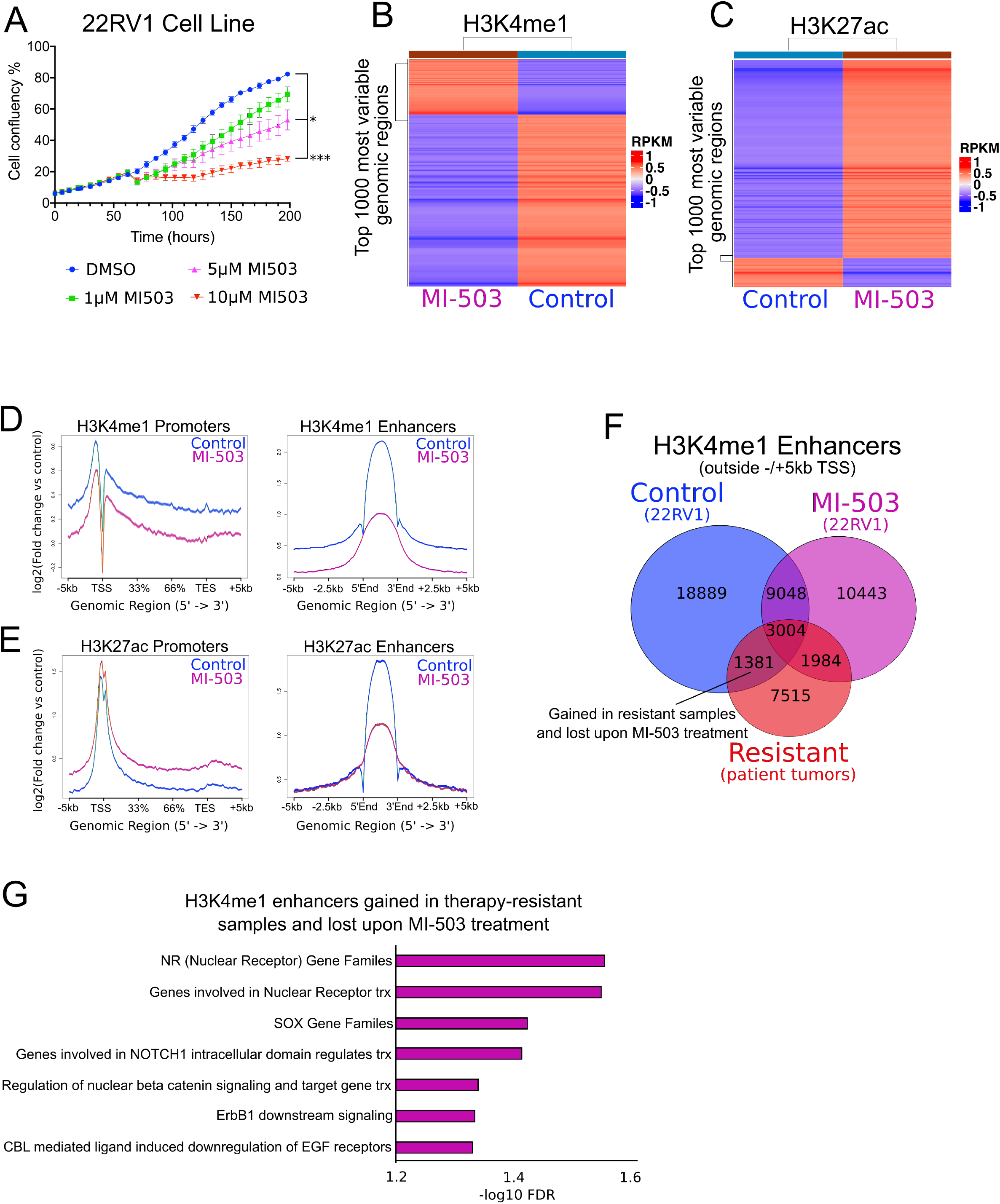
MENIN inhibition reduces H3K4me1-associated enhancers within critical resistance pathways. A) Representative proliferation assays in 22RV1 cell lines treated with DMSO (Control), enzalutamide (MV3100), menin inhibitor (MI-503) or MV3100 + MI-503 inhibitor combinations for 48 hours. n = minimum of 3 biological and 3 technical replicates in each condition. P-values based on multiple comparison ANOVA between groups. * = p < 0.05, *** = p < 0.0001 B-C) Heatmap displaying the top 1000 most variable genomic regions for H3K4me1 and C) H3K27ac RPKM values in 22RV1 cell lines treated with DMSO (Control) or 10μM MI-503 for 48 hours. D-E) Average density profiles displaying promoter (within −/+5kbTSS) and enhancer (outside −/+5kbTSS) regions for H3K4me1 and E) H3K27ac in 22RV1 cell lines treated with DMSO (Control) or 10μM MI-503 for 48 hours. F) Venn diagram displaying overlapping H3K4me1 enhancer regions gained in resistant tumors and 22RV1 cell lines treated with DMSO (Control) or 10μM MI-503 for 48 hours. G) Significant GREAT MSigDB Pathways and HGNC Gene Families based on H3K4me1 enhancers gained in therapy-resistant tumors and Control 22RV1 cell, and lost upon 10μM MI-503 treatment from F. Hyper FDR q-values are displayed as −log10.

### H3K4me1-associated enhancers are enriched for self-regulatory transcription factors in therapy resistant prostate cancers

Enhancers are epigenetic modules that regulate cell-type specific transcriptional programs by providing platforms for binding of key TFs that act as drivers of the associated phenotypes. Hence, we defined “core TFs” that likely act as key regulators of the therapy resistant enhancer signature by employing a previously published logic (Saint-Andre et al., 2016) based on core TF identification in iPSCs/stem cells (such as OCT4, SOX2, KLF4 and NANOG). Accordingly, these TFs must satisfy the following criteria (**Figure 4A)**: 1) TF binding motifs should be enriched in the enhancers specific to those clusters, 2) TF should be highly expressed in the therapy resistant tumors, and 3) TF should self-regulate themselves by enhancer activation in its own vicinity for self-regulation. Hence, we identified and integrated datasets in the following steps; 1) determine resistant-specific H3K4me1 enhancer regions, 2) predict TFs whose binding sites are enriched in therapy resistant enhancers, and 3) determine the expression level of predicted TF gene within gained treatment-specific enhancers (**Figure 4A**). Within these gained enhancers we identified motifs for core TFs known to drive CRPC or critical developmental programs. These included well known prostate cancer drivers such as AR (Malik et al., 2015; Takeda et al., 2018), FOXA1 (Parolia et al., 2019; Yang and Yu, 2015) and HOXB13 (Ewing et al., 2012; Kron et al., 2017), as well as developmental TFs associated with embryogenesis and stem cell pluripotency, such as SOX factors (Sarkar and Hochedlinger, 2013) (**Figure 4B, 4C** and **Table S2**). Importantly, these TFs are expressed in our dataset (assessed by H3K79me2 as a proxy for RNA expression) or in a large number of patient samples treated with Abiraterone-Enzalutamide (ABI-ENZ) hormone therapy (≥20 FPKM median expression) from the SU2C-PCF dataset (Abida et al., 2019) (**Figure 4D, S6B** and **Table S3**). On a genome-wide level, we found that the same canonical TFs (AR, FOXA1 and HOXB13) occupy the resistant H3K4me1 enhancers (**Figure 4E-4F).** Interestingly, these co-occupied enhancers are associated with pathways known to drive various prostate developmental programs (Pomerantz et al., 2015) (**Figure 4G-H** and **Table S4**), demonstrating a connection between H3K4me1-associated enhancer activation and the core TFs that may drive therapy resistance. Consistently, we also observed novel and previously identified developmental intergenic enhancers (Takeda et al., 2018) within *AR, FOXA1* and *HOX13* loci that were further reduced by MI-503 treatment in 22RV1 cells (**Figure 4I-J**). Notably, MI-503 targetable elements within the *SOX4* locus resemble those of developmental enhancers, displaying binding of AR, FOXA1 and HOXB13 (**Figure 4I-J**), further implicating its importance as potential member in a core TF network responsible for therapy-resistance.

**Figure 4:**
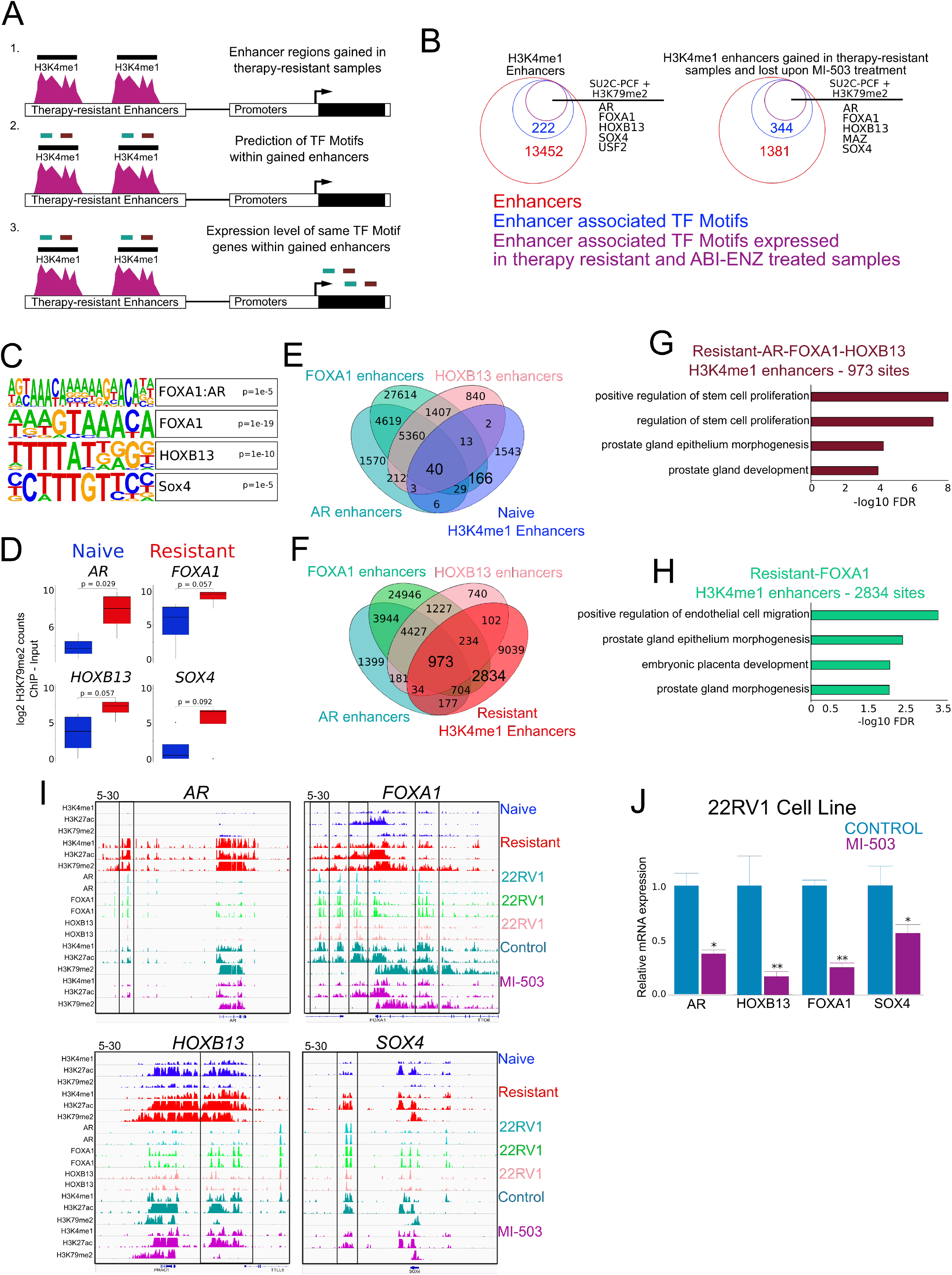
H3K4me1-associated enhancers are enriched for self-regulatory developmental transcription factors in therapy-resistant prostate cancers. A) Method used to identify core TF network in therapy-resistant prostate cancer samples; 1) determine resistant-specific H3K4me1 and active (H3K4me1+H3K27ac) enhancer regions 2) Predict TF binding sites by searching enhancers for DNA sequence TF motifs 3) Determine the expression level of predicted TF gene with gained treatment-specific enhancers B) Venn diagram displaying the total number of enhancers (outside −/+5kbTSS) in resistant samples for H3K4me1 and active regions, the number of TF motifs identified within these enhancers, and the number of TF genes expressed by H3K79me2 or within patient tumor samples treated with ABI-ENZ (≥20 FPKM) from the SU2C-PCF dataset (Abida et al., 2019). C) Motifs identified in H3K4me1 or active enhancers gained in resistant tumor samples profiled in this study. We limited this analysis to those enhancers with high expression (assayed by H3K79me2) and in patient tumor samples treated with ABI-ENZ (≥20 FPKM) from the SU2C-PCF dataset (Abida et al., 2019). D) ChIP-seq count data (log2) displaying H3K79me2 for TF identified in enhancers gained in resistant tumor samples that are also expressed within patient tumor samples treated with ABI-ENZ (≥20 FPKM) from the SU2C-PCF dataset (Abida et al., 2019). E-F) Venn diagram analysis displaying active enhancers gained in naive and F) resistant tumors with AR, FOXA1 and HOXB13 TF from 22RV1 cells (Kron et al., 2017). G-H) Significant GREAT GO Biological Process based on resistant active enhancers overlapping AR, FOXA1 and HOXB13 or H) FOXA1 enhancers in 22RV1 cells. I) ChIP-seq tracks displaying H3K4me1, H3K27ac and H3K79me2 in naive and resistant tumors, the AR, FOXA1 and HOXB13 TF from WT 22RV1 cells (Kron et al., 2017), and H3K4me1, H3K27ac and H3K79me2 in WT 22RV1 cells treated with DMSO (Control) or 10μM MI-503 on the *AR, FOXA1, HOXB13, SOX4* and *LEF1* locus. Black bar denotes putative H3K4me1-associated developmental enhancers that are lost upon 10μM MI-503 treatment. J) Associated RT-qPCR analysis for *AR, HOXB13, FOXA1* and *SOX4* mRNA expression levels in 22RV1 cells treated with DMSO (Control) or 10μM MI-503 for 48h. *GAPDH* was used as a housekeeping gene. P-values represent t-test comparisons between control and MI-503 treatment. * = p < 0.05, ** = p < 0.005. n = minimum of 2 biological and 3 technical replicates in each condition.

### SOX4 is critical for proliferation of enzalutamide-resistant cells and disrupts the H3K4me1-marked enhancer landscape

We hypothesized that as a core TF, SOX4 should be critical for growth of enzalutamide-resistant prostate cancer cells and be able to reprogram the enhancer landscape, specifically those associated with therapy resistance. Therefore, we knocked down *SOX4* levels in 22RV1 cells (**Figure 5A**). Consistent with our hypothesis and as shown in previous studies (Liu et al), we noted that SOX4 knock down reduced proliferation and 2D colony formation (**Figure 5B-5C),** yet these cells did not show any further loss of proliferation when treated with enzalutamide (**Figure S6C**). Since enhancers function as TF binding platforms and we identified SOX4 motifs in H3K4me1-associated elements, we next determined whether the knock-down of *SOX4* could influence the enhancer landscape. Interestingly, cells harboring two SOX4 shRNAs showed loss of H3K4me1-associated enhancer peaks that are specifically gained in resistant tumors (**Figure 5D**).

**Figure 5:**
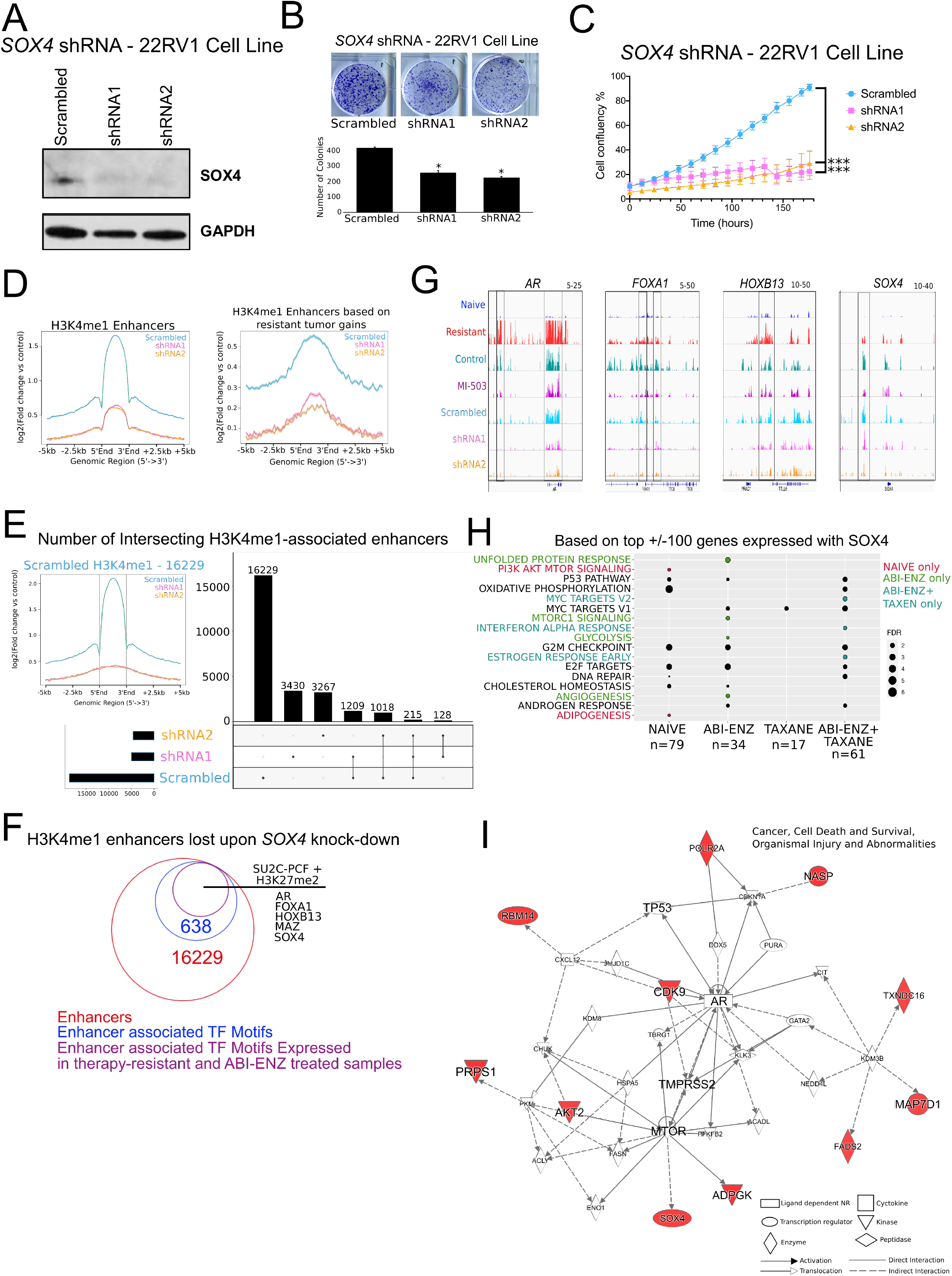
Loss of SOX4 reduces cell growth and decreases H3K4me1-associated loci. A) Western blotting analysis for *SOX4* protein expression levels in Control (Scrambled) or *SOX4* knock-down (shRNA-A1 and A2) 22RV1 cell lines. GAPDH protein expression levels were used as a loading control. B) 2d proliferation assays in Control (Scrambled) or *SOX4* shRNA (shRNA-A1 and A2) 22RV1 cell lines. P-values represent t-test comparisons between control and each *SOX4* shRNA based on number of total colonies. * = p < 0.05. Representative experiment shown. n = minimum of 2 biological and 2 technical replicates in each condition. C) Proliferation assays representing confluence percentage in Control (Scrambled) or *SOX4* shRNA (shRNA-A1 and A2) 22RV1 cell lines. P-values based on multiple comparison ANOVA between control and *SOX4* shRNA. *** = p < 0.0001. n = minimum of 2 biological and 4 technical replicates in each condition. D) Average density profiles displaying enhancer regions (outside −/+5kbTSS) of H3K4me1 in Control (Scrambled) or *SOX4* shRNA (shRNA-A1 and A2) 22RV1 cell lines (Left). Average density profiles displaying promoter and enhancer regions of H3K4me1 in Control (Scrambled) or *SOX4* shRNA (shRNA-A1 and A2) 22RV1 cell lines based on H3K4me1 gains in resistant tumor samples. E) Upset plot displaying intersecting H3K4me1-associated enhancers in Control (Scrambled) and *SOX4* shRNA (shRNA-A1 and A2) 22RV1 cell lines. F) Venn diagram displaying Control (Scrambled) H3K4me1 enhancers, the number of TF motifs identified within these enhancers, and the number of TF genes expressed by H3K79me2 or within patient tumor samples treated with ABI-ENZ (≥20 FPKM) from the SU2C-PCF dataset (Abida et al., 2019). G) ChIP-seq tracks displaying H3K4me1 on the *AR, FOXA1, HOXB13* and *SOX4* loci in naive and resistant tumors, 22RV1 lines treated with DMSO (Control) or 10μM MI-503, and 22RV1 lines expressing Control (Scrambled) or *SOX4* shRNA (shRNA-A1 and A2). H) Top 10 significant HALLMARK pathways from GSEA/MSigDB based on the top −/+100 genes co-expressed with *SOX4* in NAÏVE, ABI-ENZ, TAXANE or ABI-ENZ+TAXANE treated tumor samples from the the SU2C-PCF dataset excluding neuroendocrine tumors. Color indicates pathway uref-lnique to each treatment group. n = number of tumors in each treatment group. I) IPA gene network analysis based on the top −/+100 genes co-expressed with *SOX4* in ABI-ENZ treated tumor samples from the the SU2C-PCF dataset (Abida et al., 2019). Legend specifies potential gene relationships. Red denotes genes directly expressed together with *SOX4*.

Determination of unique enhancer regions present in control (Scrambled) cells and lost in either *SOX4*-shRNA (A1 and A2) identified a total of 16,229 H3K4me1-associated elements (**Figure 5E**). Importantly, within these elements, we identified TF motifs for the core CRPC drivers *AR*, *FOXA1, HOXB13* as well as *SOX4* (**Figure 5F** and **Table S5** and **S6**), which displayed a decrease of H3K4me1 similar to that of MI-503 treatment (**Figure 5G**). The loss of these H3K4me1 enhancers, as well as genes co-expressed together with *SOX4* (top +/− 100 genes) upon ABI-ENZ exposure, were enriched for “Androgen Response” and “Androgen Receptor Activity” (**Figure 5H, S6D** and **Table S7** and **S8**). This was also observed using gene network analysis which suggests *SOX4* expression is associated with critical therapy-resistant drivers upon ABI-ENZ exposure, including *AR*, *TMPRSS2* and *TP53* (**Figure 5I** and **S6E**). With the ability of *SOX4* to inhibit cell growth and disrupt the H3K4me1-associated enhancer landscape these results suggest SOX4 may function as one of the TF necessary to stabilize the enhancer landscape during the acquisition of therapy-resistance.

## DISCUSSION

Emerging evidence has suggested the misregulation of non-coding enhancer elements may play a key role in the progression of localized PC to CRPC. In primary prostate tumors, H3K27ac profiling identified distinct cis-regulatory landscapes between TMPRSS2-ERG-positive and TMPRSS2-ERG-negative samples (Kron et al., 2017). In a subset of these patients, the overexpression of ERG co-opts FOXA1 and HOXB13 to promote an active H3K27ac chromatin landscape associated with increased migration and invasion. On the *AR* locus, a somatically acquired, functional H3K27ac-associated enhancer has been identified as a driver of *AR* expression in CRPC (Takeda et al., 2018). This enhancer also displayed a pattern of epigenetic marks consistent with a developmental enhancer including the co-occupancy of key TFs FOXA1 and HOXB13. Taken together, these previous studies give rise to the notion the progression of PC to CRPC is governed by an interconnected TF network that includes misappropriate enhancer activation and the aberrant binding pattern of factors such as AR, FOXA1 and HOXB13.

Using chromatin state profiling encompassing six core histone modifications, we expand on these findings demonstrating that H3K4me1-associated active enhancers are enriched in therapy-resistant prostate tumor samples. Differential binding, average density and identification of resistant-specific sites revealed H3K4me1-associated loci were enriched in promoter and enhancer regions. While we did not observe enrichment of H3K27ac specific to resistant samples, a large number of H3K4me1 enhancers displayed co-occupancy with H3K27ac (3176 sites) and were associated with pathways known to drive PC progression and development, thus providing a key epigenomic mechanism that may contribute to therapy resistance. Since H3K4me1 often supersedes nucleosomal depletion and H3K27ac enhancer activation (Calo and Wysocka, 2013), and with over 10,000 H3K4me1-marked enhancers not associated with H3K27ac, further investigation is needed to determine whether these elements are necessary for H3K27ac recruitment or associated with other active modifications such as H3K9ac and H3K18ac.

In CRPC, compensatory mechanisms can lead to persistent AR expression and activity after androgen ablation or anti-androgen treatment (Chen et al., 2004; Culig and Santer, 2012). CRISPRi against the *AR* enhancer element showed deacetylation of this region effectively suppresses AR signaling and decreased sensitivity to enzalutamide (Takeda et al., 2018). Small molecule inhibition of MENIN, through an interaction with the MLL complex, blocks AR signaling and the growth of castration resistant tumors in mice (Malik et al., 2015). MLL is a component of a SET-1-like histone methyl-transferase complex that holds intrinsic H3K4 methyl-transferase activity (Dou and Hess, 2008). MENIN is necessary for MLL target gene expression (Hughes et al., 2004) and its increased expression correlates with poor patient survival. Clinical trials targeting MENIN-MLL (KO-539; NCT04067336) are currently underway in patients with refractory or relapsed Acute Myeloid Leukemia who do not respond or are ineligible for standard therapies. Using MENIN inhibition (MI-503) we found that blocking MLL function was able reduce proliferation and disrupt H3K4me1-associated enhancers in 22RV1 enzalutamide-resistant cells. The association of H3K4me1 with various nuclear receptor and developmental pathways (i.e. AR, SOX and NOTCH), and the ability of MENIN inhibition to disrupt functional enhancers regulating genes known to drive therapy-resistance, such as *AR* and *HOXB13*, identifies a plausible mechanism for therapeutically targeting advanced, therapy-resistant prostate tumors by blocking H3K4me1-marked enhancer elements that cannot be targeted by traditional hormone ablation therapies.

We found that therapy-resistant prostate cancers may be maintained by a network of developmental TFs which control their own gene expression by binding within H3K4me1 or active enhancers. We observed well-known (AR, FOXA1 and HOXB13) and less described (SOX4) developmental TFs are enriched within *de novo* H3K4me1-marked enhancers, expressed in resistant tumors used in this study or published patient tumors treated with ABI-ENZ (Abida et al., 2019), and these enhancers associate with the same TF gene loci. This process is similar to that of key regulators of ESCs, in which a small group of self-regulatory TF dominates control of a large gene expression network necessary for pluripotency, self-renewal and tissue-specific differentiation (Kim et al., 2008; Takahashi and Yamanaka, 2006). The ability of master or pioneer TFs (such as AR, FOX, HOX, and SOX) to alter chromatin architecture and regulate the expression of various gene subsets, including their own, further suggests a mechanism in which therapy-resistant prostate cancers are governed by a self-regulatory TF-network that promotes therapy-resistance. Since H3K4me1 generally precedes enhancer activation, and all forms of H3K4 methylation interfere with DNA methylation (i.e. Dnmt3L binding) (Calo and Wysocka, 2013), it is plausible that H3K4me1 functions to establish a chromatin environment necessary for the binding of core TF and subsequent enhancer activation in therapy-resistant prostate cancers.

Gene expression studies have identified increased *SOX4* expression in a variety of cancers and it is recognized as one of the 64 "cancer signature" genes (Cancer Genome Atlas Research, 2014; Liu et al., 2006; Rhodes et al., 2004; Vervoort et al., 2013a; Vervoort et al., 2013b). In PC, SOX4 expression is essential for tumorigenesis (Bilir et al., 2016) and its upregulation is associated with aggressive tumors and poor prognosis in patients (Liu et al., 2006). Here, we demonstrate that SOX4 may promote proliferation in therapy-resistant cells by functioning as one of the TF involved in establishing the H3K4me1-associated enhancer landscape. A primary role of enhancer elements is their ability to function as TF binding platforms (Calo and Wysocka, 2013). Generally, this activation requires multiple TFs in order to ensure the activation of lineage-specific genes by facilitating particular accessory factors to the promoter region. The identification of SOX4 binding sites within H3K4me1-associated enhancers together with other canonical TF (such as AR, FOXA1 and HOXB13), and the loss of H3K4me1 enhancers upon *SOX4* knock-down, together suggests it may function as one of the scaffolding TF necessary to stabilize the enhancer landscape in therapy-resistant prostate cancers. Importantly however, the moderate influence *SOX4* knock-down has on cell growth further suggests that the loss of a single TF, with the potential exception of AR or FOXA1, is not enough to sensitize resistant cells to enzalutamide. Further investigation is needed to determine the direct link between SOX4 and AR, FOXA1 and HOXB13 gene regulatory functions during acquisition of therapy resistance, and whether this process occurs in a larger cohort of patient samples. Overall, we identify reprogramming of H3K4me1-marked enhancers as an epigenetic feature associated with therapy-resistance and further suggest the use of MLL-complex inhibitors as a potential therapeutic strategy in advanced prostate cancers that do not respond to traditional therapy.

## Supporting information

GREAT pathways based on H3K4me1 enhancers in resistant tumors and Control 22RV1 cells lost upon MENIN treatment

H3K4me1 and active enhancer TF motifs in resistant tumors and Control 22RV1 cells lost upon MENIN treatment

H3K4me1 and active enhancer TF motifs in resistant tumors and Control 22RV1 cells lost upon MENIN treatment with H3K79me2 counts and SU2C-PCF FPKM

GREAT pathways based on H3K4me1 enhancers in resistant tumors overlapping AR, FOXA1 and HOXB13 in 22RV1 cells

Scrambled H3K4me1 enhancer TF motifs in 22RV1 cells

Scrambled H3K4me1 enhancer TF motifs in 22RV1 cells with H3K79me2 counts and SU2C-PCF FPKM

GSEA pathways based on the 200 genes expressed with SOX4 in ABI-ENZ treated tumor samples from the SU2C-PCF dataset

Top 200 genes expressed with SOX4 in in ABI-ENZ treated tumor samples from the SU2C-PCF dataset

## ACKNOWLEDGEMENTS

We are grateful to Kadir C. Akdemir, Yonathan Lissanu Deribe, Anand Singh, Scott Callahan, Veena Kochat, Sharon Landers, Angela Bhalla and Keila Torres for helpful discussions and proof reading the manuscript. The work and people were supported by grants from the National Institutes of Health (CA160578, CA222214 to K.R; CA157919 and CA016672 to SMF Core), and MD Anderson Cancer Center (Prostate Cancer Moon Shots and Start-up funds to K.R.; Center for Cancer Epigenetics Fellowship to C.J.T.).

## AUTHOR CONTRIBUTIONS

CJT and KR conceptualized and designed the study; MM, CJT, and NEA generated the ChIP-seq data; CJT, MM, MT and SBA processed ChIP-seq data; CJT, MT and ATR performed data analysis; CJT performed cell growth and proliferation assays; MM quantified proliferation assays; CJT performed SOX4-shRNA transduction experiments; AKG performed western blotting analysis; EO and JS provided technical help; CL, PT, FG, TT provided patient material and resources; CJT and KR wrote and prepared the manuscript. All authors read and approved the manuscript.

## METHODS

### Lead Contact

Further information and requests for resources and reagents should be directed to and will be fulfilled by the Lead Contact, Kunal Rai.

### Data and code availability

Publicly available RNA-seq data used in this study can be downloaded from the NCBI GEO BioProject database with the following accession number GSE3325 and from cBioPortal https://www.cbioportal.org/study/summary?id=prad_su2c_2015 or Github/cBioPortal https://github.com/cBioPortal/datahub/tree/master/public/prad_su2c_2019.

Code used in this study is available on https://gitlab.com/railab, https://rpubs.com/cjt5 and https://github.com/sccallahan/ChromExploreR.

### Human Subjects

Primary prostate tumors were collected at the MD Anderson Cancer Center and reviewed by a genitourinarypathologist for tumor content. Tumors with >60% content were chosen for epigenomic analyses presented here. We utilized frozen tumor samples, either flash frozen or OCT frozen, for epigenome profiling.

### Cell Lines

22RV1 cell lines were purchased from ATCC (CRL-2505) and maintained in RPMI growth media supplemented with 10% Heat-Inactivated Fetal Bovine Serum (GIBCO) and 1% Penicillin Streptomycin (HyClone). Media was changed every third day.

## METHOD DETAILS

### Chromatin immunoprecipitation assays

ChIP assays were performed as described previously (Terranova et al., 2018). Briefly, for tumor samples, ~50 mg of tissue (~8 mg per histone modification antibody) were disassociated manually in 2 mL of Hanks’ Balanced Salt Solution (HBSS) using a sterile razor blade. The tissue was minced into 3 to 4 mm pieces for approximately 5 min in a sterile tissue culture dish. The tissue was then transferred into a dissociator tube and another 8 mL of HBSS was added. For prostate tumors, the dissociator was run using the following cycles each one time in this order: h_tumor_01.01, h_tumor_02.01, h_tumor_03.01 and m_heart_02.01. For cell lines, 2 × 10^7^ cells were scraped and collected into a 15mL tube. Tissue and cells were cross-linked with 1% (wt/ vol) formaldehyde for 10 min at 37 °C with shaking. After quenching with 150 mM glycine for 10 min at 37 °C with shaking, tumors or cell lines were washed twice with ice-cold PBS and frozen at −80 °C for further processing. Cross-linked pellets were thawed and lysed on ice for 30 min in ChIP harvest buffer (12 mM Tris-Cl, 1 × PBS, 6 mM EDTA, 0.5% SDS) with protease inhibitors (Sigma). Lysed cells were sonicated with a Bioruptor (Diagenode) to obtain chromatin fragments (~200–500 bp) and centrifuged at 15,000 × g for 15 min to obtain a soluble chromatin fraction. In parallel with cellular lysis and sonication, antibodies (2ug/IP for tissue and 5 μg/3 × 106 for cell lines) were coupled with 30 μl of magnetic protein G beads in binding/blocking buffer (PBS + 0.1% Tween + 0.2% BSA) for 2 h at 4 °C with rotation. Soluble chromatin was diluted five times using ChIP dilution buffer (10 mM Tris-Cl, 140 mM NaCl, 0.1% DOC, 1% Triton X, 1 mM EDTA) with protease inhibitors and added to the antibody-coupled beads with rotation at 4 °C overnight. Immune complexes were washed five times with cold RIPA buffer (10mM Tris–HCl, pH 8.0, 1mM EDTA, pH 8.0, 140mM NaCl, 1% Triton X-100, 0.1% SDS, 0.1% DOC), twice with cold high-salt RIPA buffer (10mM Tris–HCl, pH 8.0, 1mM EDTA, pH 8.0, 500mM NaCl, 1% Triton X-100, 0.1% SDS, 0.1% DOC), and twice with cold LiCl buffer (10mM Tris–HCl, pH 8.0, 1mM EDTA, pH 8.0, 250mM LiCl, 0.5% NP-40, 0.5% DOC). After washing, samples were treated with elution buffer (10 mM Tris-Cl, pH 8.0, 5 mM EDTA, 300 mM NaCl, 0.5% SDS), RNase A, and Proteinase K, and cross-links were reversed overnight. ChIP DNA was purified using AMPure XP beads (Agencourt) and quantified using the Qubit 2000 (Invitrogen) and TapeStation4200 (Agilent). Libraries for Illumina sequencing were generated following the New England BioLabs (NEB) Next Ultra DNA Library Prep Kit protocol. A total of 10 cycles were used during PCR amplification for the generation of all ChIP-seq libraries. Amplified ChIP DNA was purified using double-sided AMPure XP to retain fragments (~200–500 bp) and quantified using the Qubit 2000 and Bioanalyzer 1000 before multiplexing.

### ChIP-seq data processing

Raw fastq reads for all ChIP-seq experiments were processed using a snakemake based pipeline https://github.com/crazyhottommy/pyflow-ChIPseq (Blecher-Gonen et al., 2013). Briefly, raw reads were first processed using FastQC (http://www.bioinformatics.babraham.ac.uk/projects/fastqc/) and uniquely mapped reads were aligned to the hg19 reference genome using Bowtie version 1.1.2 (Langmead et al., 2009). Duplicate reads were removed using SAMBLASTER (Faust and Hall, 2014) before compression to bam files. To directly compare ChIP-seq samples uniquely mapped reads for each mark were downsampled per condition to 10 million, sorted and indexed using samtools version 1.5 (Li et al., 2009). To visualize ChIP-seq libraries on the IGV genome browser, we used deepTools version 2.4.0 (Ramirez et al., 2016) to generate bigWig files by scaling the bam files to reads per kilobase per million (RPKM). Super ChIP-seq tracks were generated by merging bam files from naive or resistant tumors, sorting and indexing using samtools and scaling to RPKM using deepTools.

### Chromatin state calls

ChromHMM (Ernst and Kellis, 2012) was used to identify combinatorial chromatin state patterns based on the histone modifications studied. Normalized bam files were converted to bed files and binarized at a 1000bp resolution using the BinarizeBed command. We specified that ChromHMM should learn a model based on 15 chromatin states. As we considered models between 8 and 30 chromatin states, we chose a 15-state model because it is large enough to identify important functional elements while still being small enough to interpret easily. To determine chromatin state differences between different groups we used a two-step process. First, using the segmentation calls from the ChromHMM output the entire genome is divided into non-overlapping windows of 10 Kb. We next count the number of times a chromatin state is observed in each of the 10 Kb windows and obtain a frequency matrix for each state in the ChromHMM model (E1-E15). In the second step, low variable genomic loci are removed from the frequency matrix and significant differences between two groups of samples types are calculated by using a nonparametric Mann Whitney Wilcoxon test with a p-value < 0.05 for each state separately.

### Identification and visualization of ChIP-seq binding sites

We used Model-based analysis of ChIP-seq (MACS) version 1.4.2 (Zhang et al., 2008) peak calling algorithm to identify H3K4me1 and H3K27ac (p-value of 1e-5) enrichment over “input” background. Active enhancers were identified by overlapping H3K4me1 and H3K27ac by a minimum of 1bp using intersectBed (Quinlan and Hall, 2010). Final peaksets used for downstream analysis were generated using mergeBed. Unique peaks for enhancers and active enhancers were identified using the concatenate, cluster and subtract tools from the Galaxy/Cistrome web-based platform (Liu et al., 2011). Unique peak identification for 22RV1 Control (DMSO) and MI-503 enhancer overlap with resistant active enhancers, 22RV1 core TF (AR, FOXA1 and HOXB13) enhancer overlap with resistant active enhancers, and 22RV1 Scrambled enhancer overlap with SOX4-shRNA enhancers were performed using Intervene (Khan and Mathelier, 2017).

### Differential Binding Analysis

Differential binding analysis for H3K4me1, H3K27ac and H3K79me2 was performed using DiffBind(Ross-Innes et al., 2012). Count data was obtained from BAM and corresponding peak files using the dba.count command with a minimum overlap = 2, score = DBA_SCORE_READS_MINUS and fragment size = 200. Differential analysis on count data was performed using DESEQ2 in the DiffBind package.

### Assigning binding sites to genes

A list of Ensembl genes was obtained from the UCSC Table browser (http://genome.ucsc.edu/). Proximal promoters were defined as ±5kb from the transcription start site (TSS). Peaks were assigned to promoters if they overlapped the promoter by a minimum of 1bp using intersectBed. Enhancers were defined as all peaks outside of the promoter region (±5kb TSS). Enhancers were identified by subtracting all peaks by proximal promoter peaks using subtractBed by a minimum of 1bp. For identification of self-regulatory TFs, enhancers peaks were assigned to genes if they were located within −/+50kb of an enhancer peak. For H3K79me2, peaks were assigned to genes if they overlapped within −5kbTSS to the Transcription End Site before transformation to count data with DiffBind. Gene body heatmaps and average density plots were generated using ngs.plot (Shen et al., 2014).

### Gene set enrichment analysis

Pathways associated with H3K4me1 and active enhancers were identified used Genomic Regions of Enrichment Annotation Tool (GREAT) with a basal setting of −/+5kb from the TSS with a 1MB extension. Control (Scrambled) H3K4me1 enhancer pathways were identified with the single gene setting with a 1MB extension. All pathways are significant based Hyper FDR q-values. For RNA-seq data we used GSEA/MSigDB (Subramanian et al., 2005) with HALLMARK settings based on expression values >20 FPKM in each condition. All pathways are significant based on FDR q-values.

### Motif Analysis

Fasta sequences were obtained from bed files using Cistrome/Galaxy and all motifs were identified using the findMotifs command in homer (Heinz et al., 2010) under default settings. For background, untreated H3K4me1 enhancers were used to compare resistant H3K4me1 enhancers; concatenated shRNA3 + shRNA4 H3K4me1 enhancers were used to compare Scrambled H3K4me1 enhancers; input files (1381 sites) were scrambled and used to compare H3K4me1 enhancers gained in therapy-resistant prostate cancers and lost upon MI-503 treatment.

### Stable Cell Line Generation and western blotting analysis

For lenti packaging, *SOX4*-shRNA (5μg) and *Scrambled*-shRNA (5μg) in a pLKO.1 puro vector was transfected together with packaging plasmids (5μg) in HEK293T cells using Lipofectamine 3000 according to manufacturer’s instructions. For transduction, lentivirus with RPMI media + 10μg of polybrene was reversed transduced with 100,000 cells of 22RV1 cells and transgene selection was performed using 3μg of puromycin. Stable cell lines were washed with ice-cold PBS then lysed for 30 minutes at 4°C with agitation in RIPA buffer (Boston BioProducts; BP-115) supplemented with a Roche Complete Mini EDTA-free protease inhibitor cocktail (Millipore Sigma; 11836170001). Lysate was centrifuged at 4°C for 15 minutes at 15,000rpm. The pellet was discarded, and protein was quantified using BCA assay (Sigma, St Louis, MO). Samples were supplemented with 2x Laemmli buffer (Bio-Rad; 1610737) and 2-mercaptoethanol, heated for 5 min at 95°C and loaded on 4-20% Criterion TGX Stain-Free gels (Bio-Rad; 5678094). Proteins were transferred to a PVDF membrane which was then blocked in 5% non-fat dry milk in 1X TBST and incubated with primary antibody overnight at 4°C in the same buffer. Protein expression was examined by western blot using anti-SOX4 (EMD Millipore. AB5803; 1:1000 dilution) or anti-GAPDH (Proteintech, 60004-1-Ig; 1:30,000 dilution). Blots were then washed with 1X TBST and probed with goat anti-rabbit IgG-HRP (CST; 7074S) or horse anti-mouse IgG-HRP (CST; 7076S). Following subsequent washing, reactive bands were detected by Amersham ECL Western Blotting Detection Reagent (RPN2106, GE Healthcare) or SuperSignal West Dura Extended Duration substrate (Thermo; 34075).

### Proliferation assays

Cell proliferation was measured and quantified using IncuCyte ZOOM system (Essen Biosciences). Briefly, 22RV1 cells were seeded at a density of 5,000 cells per well in 96-well plates. Cells were treated with designated concentration of MDV-3100 (S1250-5mg) or MI-503 (50-136-5300) in DMSO and media was replaced every third day. Cell proliferation is plotted as confluence percentage. For *SOX4* shRNA assays cells were seeded at a density of 5,000 cells per well in 96-well plates and standard media was replaced every third day. For 2d experiments, cells were seeded at a density of 3, 000 cells per well in 6-well plates. Cells were grown for a total of three weeks, washed with 1% PBS and incubated with 10% formalin for 10 minutes. Cells were washed again with 1% PBS and incubated in crystal violet staining for 10 minutes. Colonies were counted using ImageJ software. For ZOOM experiments p-values were calculated by a paired t-test for each final time point using graph pad prism. For 2d assays experiments p-values were calculated by a paired t-test for each final time point using SPSS software.

### Quantitative PCR

Total RNA was isolated using RNeasy Mini Kit (QIAGEN) and 1μg of RNA from each sample was reverse transcribed using SuperScript VILO cDNA Synthesis Kit (Thermo Fisher). qPCR was performed with 2μl of undiluted cDNA in triplicate for each primer set. *GAPDH* was used as a housekeeping gene and data was plotted and quantified using https://doi.org/10.7717/peerj.4473

## REFERENCES

Abida, W., Cyrta, J., Heller, G., Prandi, D., Armenia, J., Coleman, I., Cieslik, M., Benelli, M., Robinson, D., Van Allen, E. M., et al. (2019). Genomic correlates of clinical outcome in advanced prostate cancer. Proc Natl Acad Sci U S A 116, 11428–11436.

Adamo, P., and Ladomery, M.R. (2016). The oncogene ERG: a key factor in prostate cancer. Oncogene 35, 403–414.

Bilir, B., Osunkoya, A.O., Wiles, W.G.t., Sannigrahi, S., Lefebvre, V., Metzger, D., Spyropoulos, D.D., Martin, W.D., and Moreno, C.S. (2016). SOX4 Is Essential for Prostate Tumorigenesis Initiated by PTEN Ablation. Cancer Res 76, 1112–1121.

Blecher-Gonen, R., Barnett-Itzhaki, Z., Jaitin, D., Amann-Zalcenstein, D., Lara-Astiaso, D., and Amit, I. (2013). High-throughput chromatin immunoprecipitation for genome-wide mapping of in vivo protein-DNA interactions and epigenomic states. Nature protocols 8, 539–554.

Brooke, G.N., and Bevan, C.L. (2009). The role of androgen receptor mutations in prostate cancer progression. Curr Genomics 10, 18–25.

Calo, E., and Wysocka, J. (2013). Modification of enhancer chromatin: what, how, and why? Mol Cell 49, 825–837.

Cancer Genome Atlas Research, N. (2014). Comprehensive molecular characterization of urothelial bladder carcinoma. Nature 507, 315–322.

Chang, K.H., Li, R., Kuri, B., Lotan, Y., Roehrborn, C.G., Liu, J., Vessella, R., Nelson, P.S., Kapur, P., Guo, X., et al. (2013). A gain-of-function mutation in DHT synthesis in castration-resistant prostate cancer. Cell 154, 1074–1084.

Chen, C.D., Welsbie, D.S., Tran, C., Baek, S.H., Chen, R., Vessella, R., Rosenfeld, M.G., and Sawyers, C.L. (2004). Molecular determinants of resistance to antiandrogen therapy. Nat Med 10, 33–39.

Culig, Z., and Santer, F.R. (2012). Androgen receptor co-activators in the regulation of cellular events in prostate cancer. World J Urol 30, 297–302.

Dou, Y., and Hess, J.L. (2008). Mechanisms of transcriptional regulation by MLL and its disruption in acute leukemia. Int J Hematol 87, 10–18.

Ernst, J., and Kellis, M. (2012). ChromHMM: automating chromatin-state discovery and characterization. Nature methods 9, 215–216.

Ewing, C.M., Ray, A.M., Lange, E.M., Zuhlke, K.A., Robbins, C.M., Tembe, W.D., Wiley, K.E., Isaacs, S.D., Johng, D., Wang, Y., et al. (2012). Germline mutations in HOXB13 and prostate-cancer risk. N Engl J Med 366, 141–149.

Faust, G.G., and Hall, I.M. (2014). SAMBLASTER: fast duplicate marking and structural variant read extraction. Bioinformatics 30, 2503–2505.

Freedland, S.J., Humphreys, E.B., Mangold, L.A., Eisenberger, M., Dorey, F.J., Walsh, P.C., and Partin, A.W. (2005). Risk of prostate cancer-specific mortality following biochemical recurrence after radical prostatectomy. JAMA 294, 433–439.

Gao, N., Ishii, K., Mirosevich, J., Kuwajima, S., Oppenheimer, S.R., Roberts, R.L., Jiang, M., Yu, X., Shappell, S.B., Caprioli, R.M., et al. (2005). Forkhead box A1 regulates prostate ductal morphogenesis and promotes epithelial cell maturation. Development 132, 3431–3443.

Gao, N., Zhang, J., Rao, M.A., Case, T.C., Mirosevich, J., Wang, Y., Jin, R., Gupta, A., Rennie, P.S., and Matusik, R.J. (2003). The role of hepatocyte nuclear factor-3 alpha (Forkhead Box A1) and androgen receptor in transcriptional regulation of prostatic genes. Mol Endocrinol 17, 1484–1507.

Heinlein, C.A., and Chang, C. (2004). Androgen receptor in prostate cancer. Endocr Rev 25, 276–308.

Heinz, S., Benner, C., Spann, N., Bertolino, E., Lin, Y.C., Laslo, P., Cheng, J.X., Murre, C., Singh, H., and Glass, C.K. (2010). Simple combinations of lineage-determining transcription factors prime cis-regulatory elements required for macrophage and B cell identities. Mol Cell 38, 576–589.

Hughes, C.M., Rozenblatt-Rosen, O., Milne, T.A., Copeland, T.D., Levine, S.S., Lee, J.C., Hayes, D.N., Shanmugam, K.S., Bhattacharjee, A., Biondi, C.A., et al. (2004). Menin associates with a trithorax family histone methyltransferase complex and with the hoxc8 locus. Mol Cell 13, 587–597.

Ishizaki, F., Nishiyama, T., Kawasaki, T., Miyashiro, Y., Hara, N., Takizawa, I., Naito, M., and Takahashi, K. (2013). Androgen deprivation promotes intratumoral synthesis of dihydrotestosterone from androgen metabolites in prostate cancer. Sci Rep 3, 1528.

Kaestner, K.H. (2010). The FoxA factors in organogenesis and differentiation. Curr Opin Genet Dev 20, 527–532.

Khan, A., and Mathelier, A. (2017). Intervene: a tool for intersection and visualization of multiple gene or genomic region sets. BMC Bioinformatics 18, 287.

Kim, J., Chu, J., Shen, X., Wang, J., and Orkin, S.H. (2008). An extended transcriptional network for pluripotency of embryonic stem cells. Cell 132, 1049–1061.

Kron, K.J., Murison, A., Zhou, S., Huang, V., Yamaguchi, T.N., Shiah, Y.J., Fraser, M., van der Kwast, T., Boutros, P.C., Bristow, R.G., et al. (2017). TMPRSS2-ERG fusion co-opts master transcription factors and activates NOTCH signaling in primary prostate cancer. Nat Genet 49, 1336–1345.

Kupelian, P.A., Mahadevan, A., Reddy, C.A., Reuther, A.M., and Klein, E.A. (2006). Use of different definitions of biochemical failure after external beam radiotherapy changes conclusions about relative treatment efficacy for localized prostate cancer. Urology 68, 593–598.

Langmead, B., Trapnell, C., Pop, M., and Salzberg, S.L. (2009). Ultrafast and memory-efficient alignment of short DNA sequences to the human genome. Genome biology 10, R25.

Lee, H.J., Hwang, M., Chattopadhyay, S., Choi, H.S., and Lee, K. (2008). Hepatocyte nuclear factor-3 alpha (HNF-3alpha) negatively regulates androgen receptor transactivation in prostate cancer cells. Biochem Biophys Res Commun 367, 481–486.

Li, J., Ng, E.K., Ng, Y.P., Wong, C.Y., Yu, J., Jin, H., Cheng, V.Y., Go, M.Y., Cheung, P.K., Ebert, M.P., et al. (2009). Identification of retinoic acid-regulated nuclear matrix-associated protein as a novel regulator of gastric cancer. Br J Cancer 101, 691–698.

Liu, P., Ramachandran, S., Ali Seyed, M., Scharer, C.D., Laycock, N., Dalton, W.B., Williams, H., Karanam, S., Datta, M.W., Jaye, D.L., et al. (2006). Sex-determining region Y box 4 is a transforming oncogene in human prostate cancer cells. Cancer Res 66, 4011–4019.

Liu, T., Ortiz, J.A., Taing, L., Meyer, C.A., Lee, B., Zhang, Y., Shin, H., Wong, S.S., Ma, J., Lei, Y., et al. (2011). Cistrome: an integrative platform for transcriptional regulation studies. Genome biology 12, R83.

Malik, R., Khan, A.P., Asangani, I.A., Cieslik, M., Prensner, J.R., Wang, X., Iyer, M.K., Jiang, X., Borkin, D., Escara-Wilke, J., et al. (2015). Targeting the MLL complex in castration-resistant prostate cancer. Nat Med 21, 344–352.

Mikkelsen, T.S., Ku, M., Jaffe, D.B., Issac, B., Lieberman, E., Giannoukos, G., Alvarez, P., Brockman, W., Kim, T.K., Koche, R.P., et al. (2007). Genome-wide maps of chromatin state in pluripotent and lineage-committed cells. Nature 448, 553–560.

Mohler, J.L. (2008). Castration-recurrent prostate cancer is not androgen-independent. Adv Exp Med Biol 617, 223–234.

Mulholland, D.J., Kobayashi, N., Ruscetti, M., Zhi, A., Tran, L.M., Huang, J., Gleave, M., and Wu, H. (2012). Pten loss and RAS/MAPK activation cooperate to promote EMT and metastasis initiated from prostate cancer stem/progenitor cells. Cancer Res 72, 1878–1889.

Parolia, A., Cieslik, M., Chu, S.C., Xiao, L., Ouchi, T., Zhang, Y., Wang, X., Vats, P., Cao, X., Pitchiaya, S., et al. (2019). Distinct structural classes of activating FOXA1 alterations in advanced prostate cancer. Nature 571, 413–418.

Pomerantz, M.M., Li, F., Takeda, D.Y., Lenci, R., Chonkar, A., Chabot, M., Cejas, P., Vazquez, F., Cook, J., Shivdasani, R.A., et al. (2015). The androgen receptor cistrome is extensively reprogrammed in human prostate tumorigenesis. Nat Genet 47, 1346–1351.

Quinlan, A.R., and Hall, I.M. (2010). BEDTools: a flexible suite of utilities for comparing genomic features. Bioinformatics 26, 841–842.

Ramirez, F., Ryan, D.P., Gruning, B., Bhardwaj, V., Kilpert, F., Richter, A.S., Heyne, S., Dundar, F., and Manke, T. (2016). deepTools2: a next generation web server for deep-sequencing data analysis. Nucleic Acids Res 44, W160–165.

Rhodes, D.R., Yu, J., Shanker, K., Deshpande, N., Varambally, R., Ghosh, D., Barrette, T., Pandey, A., and Chinnaiyan, A.M. (2004). Large-scale meta-analysis of cancer microarray data identifies common transcriptional profiles of neoplastic transformation and progression. Proc Natl Acad Sci U S A 101, 9309–9314.

Roadmap Epigenomics, C., Kundaje, A., Meuleman, W., Ernst, J., Bilenky, M., Yen, A., Heravi-Moussavi, A., Kheradpour, P., Zhang, Z., Wang, J., et al. (2015). Integrative analysis of 111 reference human epigenomes. Nature 518, 317–330.

Ross-Innes, C.S., Stark, R., Teschendorff, A.E., Holmes, K.A., Ali, H.R., Dunning, M.J., Brown, G.D., Gojis, O., Ellis, I.O., Green, A.R., et al. (2012). Differential oestrogen receptor binding is associated with clinical outcome in breast cancer. Nature 481, 389–393.

Ryan, C.J., Smith, M.R., de Bono, J. S., Molina, A., Logothetis, C.J., de Souza, P., Fizazi, K., Mainwaring, P., Piulats, J.M., Ng, S., et al. (2013). Abiraterone in metastatic prostate cancer without previous chemotherapy. N Engl J Med 368, 138–148.

Saint-Andre, V., Federation, A.J., Lin, C.Y., Abraham, B.J., Reddy, J., Lee, T.I., Bradner, J.E., and Young, R.A. (2016). Models of human core transcriptional regulatory circuitries. Genome Res 26, 385–396.

Sarkar, A., and Hochedlinger, K. (2013). The sox family of transcription factors: versatile regulators of stem and progenitor cell fate. Cell Stem Cell 12, 15–30.

Scher, H.I., Fizazi, K., Saad, F., Taplin, M.E., Sternberg, C.N., Miller, K., de Wit, R., Mulders, P., Chi, K.N., Shore, N.D., et al. (2012). Increased survival with enzalutamide in prostate cancer after chemotherapy. N Engl J Med 367, 1187–1197.

Scher, H.I., and Sawyers, C.L. (2005). Biology of progressive, castration-resistant prostate cancer: directed therapies targeting the androgen-receptor signaling axis. J Clin Oncol 23, 8253–8261.

Shen, L., Shao, N., Liu, X., and Nestler, E. (2014). ngs.plot: Quick mining and visualization of next-generation sequencing data by integrating genomic databases. BMC Genomics 15, 284.

Siegel, R., Ma, J., Zou, Z., and Jemal, A. (2014). Cancer statistics, 2014. CA Cancer J Clin 64, 9–29.

Subramanian, A., Tamayo, P., Mootha, V.K., Mukherjee, S., Ebert, B.L., Gillette, M.A., Paulovich, A., Pomeroy, S.L., Golub, T.R., Lander, E.S., et al. (2005). Gene set enrichment analysis: a knowledge-based approach for interpreting genome-wide expression profiles. Proc Natl Acad Sci U S A 102, 15545–15550.

Takahashi, K., and Yamanaka, S. (2006). Induction of pluripotent stem cells from mouse embryonic and adult fibroblast cultures by defined factors. Cell 126, 663–676.

Takeda, D.Y., Spisak, S., Seo, J.H., Bell, C., O’Connor, E., Korthauer, K., Ribli, D., Csabai, I., Solymosi, N., Szallasi, Z., et al. (2018). A Somatically Acquired Enhancer of the Androgen Receptor Is a Noncoding Driver in Advanced Prostate Cancer. Cell 174, 422–432 e413.

Terranova, C., Tang, M., Orouji, E., Maitituoheti, M., Raman, A., Amin, S., Liu, Z., and Rai, K. (2018). An Integrated Platform for Genome-wide Mapping of Chromatin States Using High-throughput ChIP-sequencing in Tumor Tissues. J Vis Exp.

Vervoort, S.J., Lourenco, A.R., van Boxtel, R., and Coffer, P.J. (2013a). SOX4 mediates TGF-beta-induced expression of mesenchymal markers during mammary cell epithelial to mesenchymal transition. PLoS One 8, e53238.

Vervoort, S.J., van Boxtel, R., and Coffer, P.J. (2013b). The role of SRY-related HMG box transcription factor 4 (SOX4) in tumorigenesis and metastasis: friend or foe? Oncogene 32, 3397–3409.

Waltering, K.K., Urbanucci, A., and Visakorpi, T. (2012). Androgen receptor (AR) aberrations in castration-resistant prostate cancer. Mol Cell Endocrinol 360, 38–43.

Yang, Y.A., and Yu, J. (2015). Current perspectives on FOXA1 regulation of androgen receptor signaling and prostate cancer. Genes Dis 2, 144–151.

Zhang, M., Behbod, F., Atkinson, R.L., Landis, M.D., Kittrell, F., Edwards, D., Medina, D., Tsimelzon, A., Hilsenbeck, S., Green, J.E., et al. (2008). Identification of tumor-initiating cells in a p53-null mouse model of breast cancer. Cancer Res 68, 4674–4682.

