## Supplemental Figures for "H3K4me1-marked Enhancer Activation in Resistant Prostate Cancers Implicates SOX4 and MENIN Inhibition as Therapeutic Strategies"

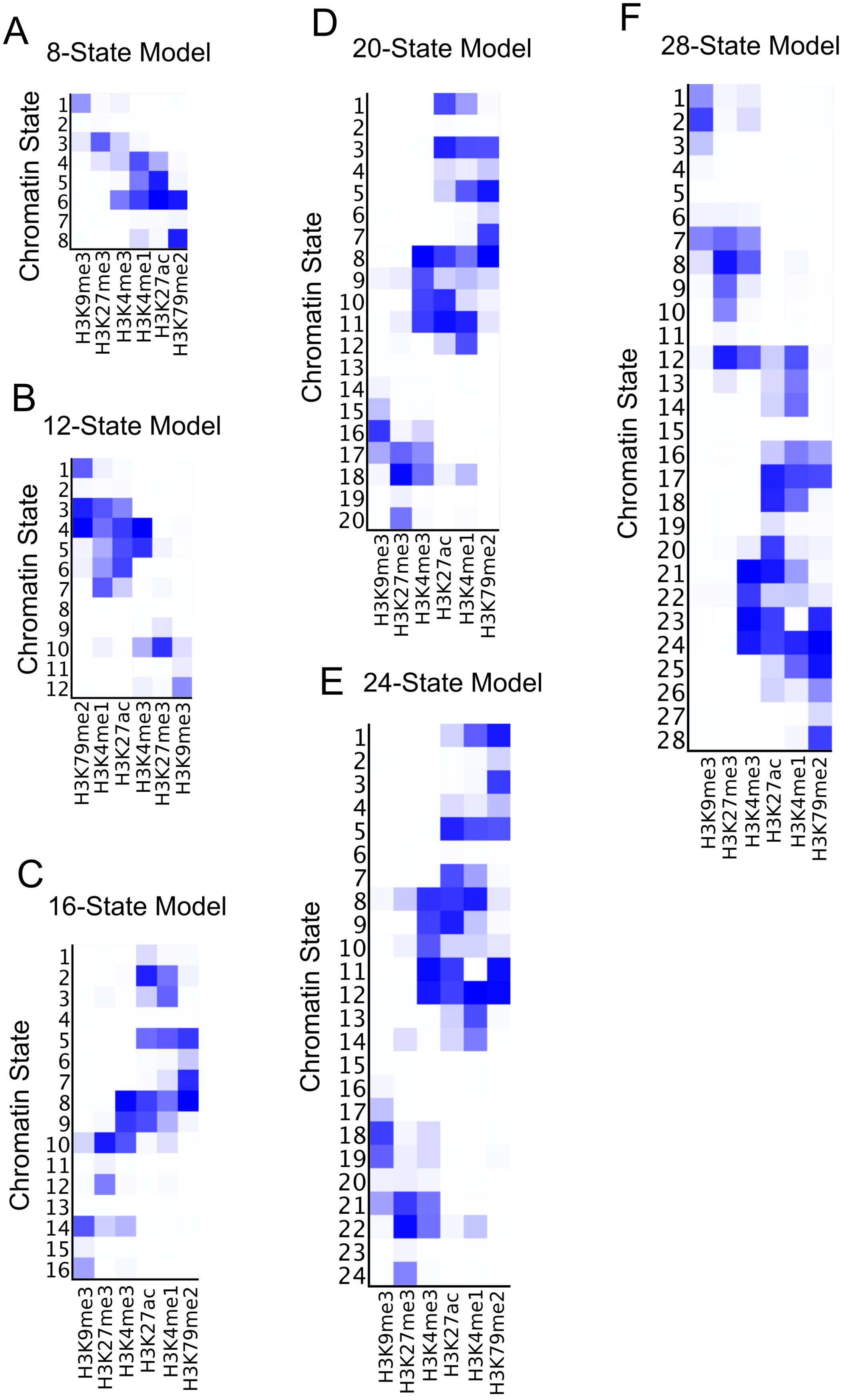


**Figure S1: Chromatin state definitions in naive and therapy-resistant prostate tumors**

A-F) Representative chromatin state definitions representing 8-, 12-, 16-, 20-, 24- and 28-state models and their associated histone mark probabilities in 4 naive and 4 resistant prostate cancer tumors.

### Figure S2

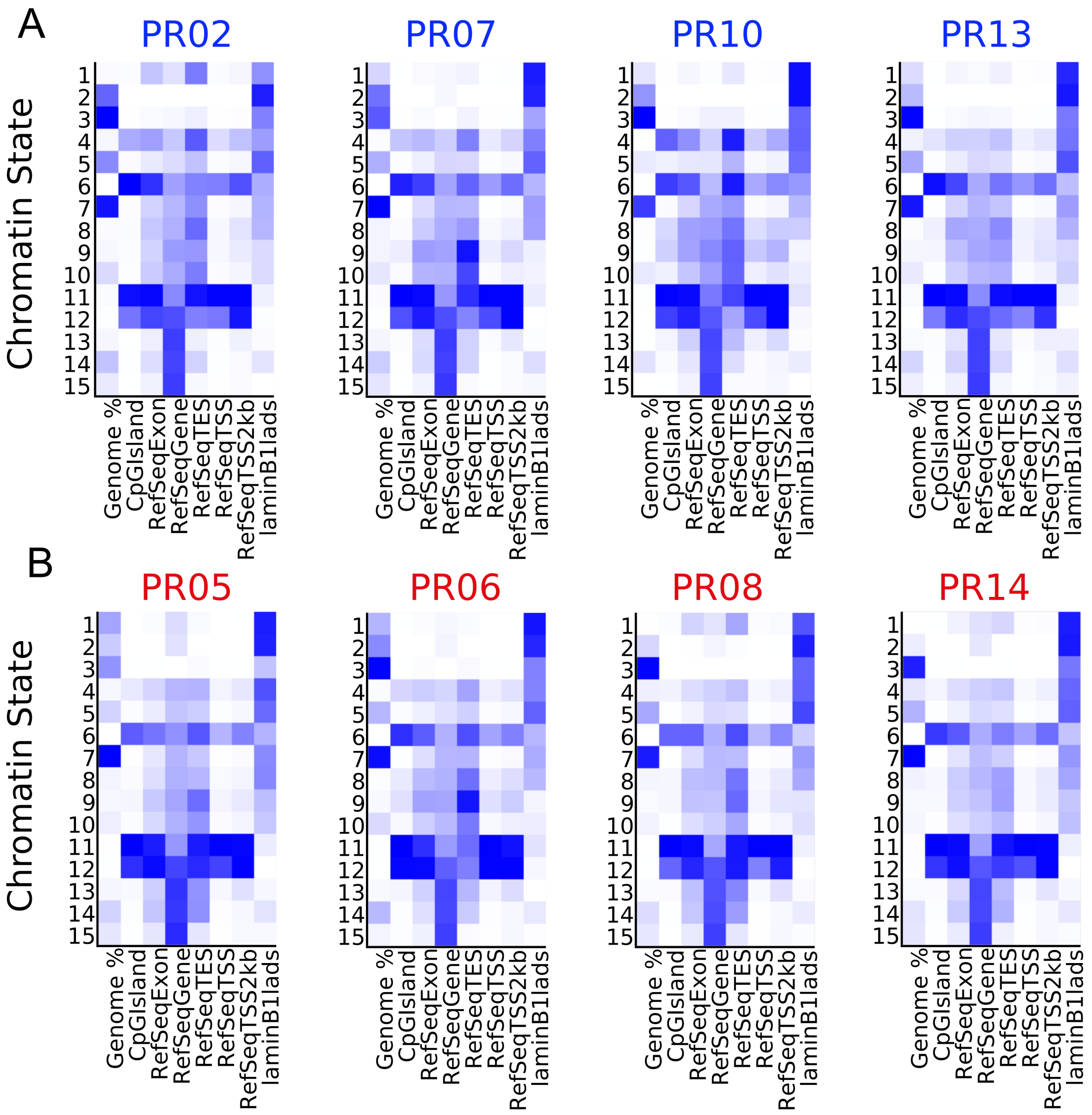


**Figure S2: Genomic enrichment definitions in naive and therapy-resistant prostate tumors**
A) Genomic enrichments for each chromatin state in 4 naive and B) 4 resistant prostate cancer tumors.

### Figure S3

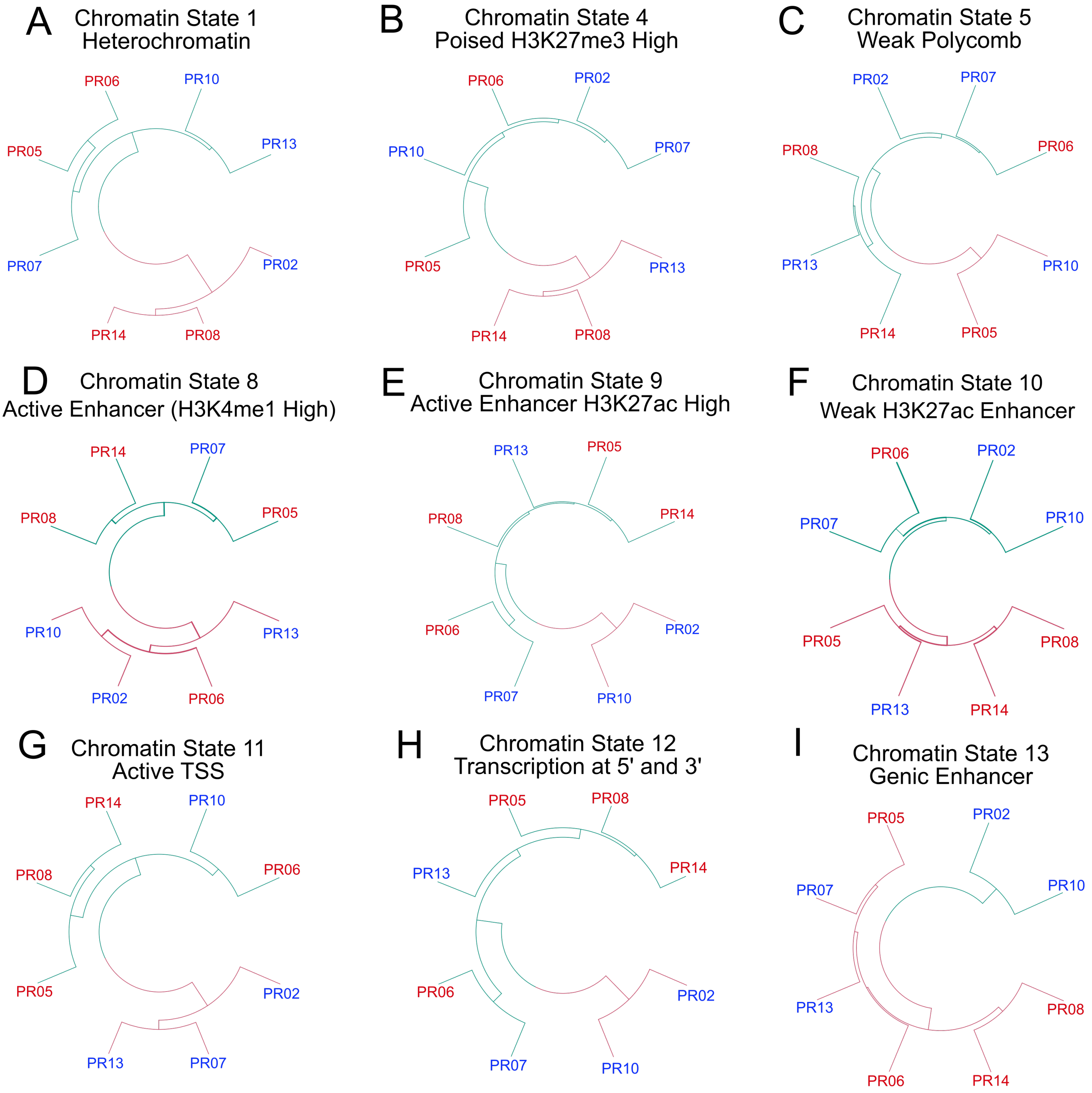


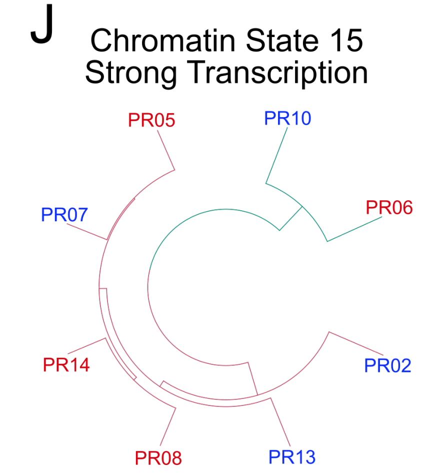


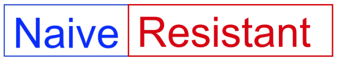


**Figure S3: Unsupervised clustering analysis of chromatin states in naive and therapy-resistant prostate tumors**
A-J) Dendrogram of each chromatin state annotated by naive or resistant tumor samples from Figure1B. This does not include weak heterochromatin (state 2), weak transcription (state 14), and low or artifact states.

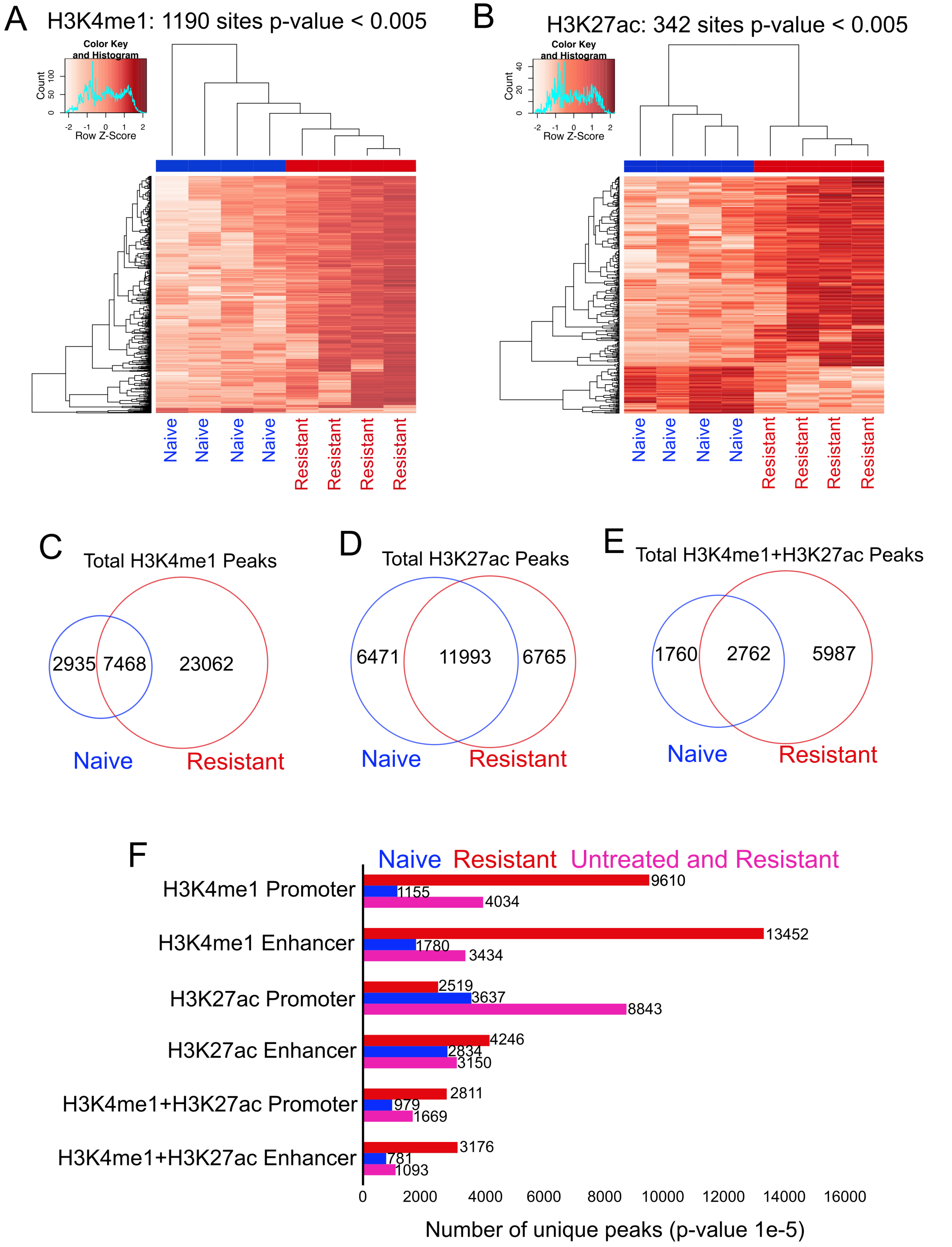


**Figure S4: H3K4me1-marked enhancers are enriched in therapy-resistant prostate tumors**
A) Heatmap displaying differential H3K4me1 and B) H3K27ac sites in naive and resistant prostate tumors. C) Venn diagram displaying total H3K4me1, D) total H3K27ac and E) total H3K4me1+H3K27ac peaks in naive and resistant prostate tumors. F) Barplot of unique H3K4me1, H3K27ac and H3K4me1+H3K27ac peaks in promoter (within -/+5kbTSS) and enhancer (outside -/+5kbTSS) genomic regions.

### Figure S5

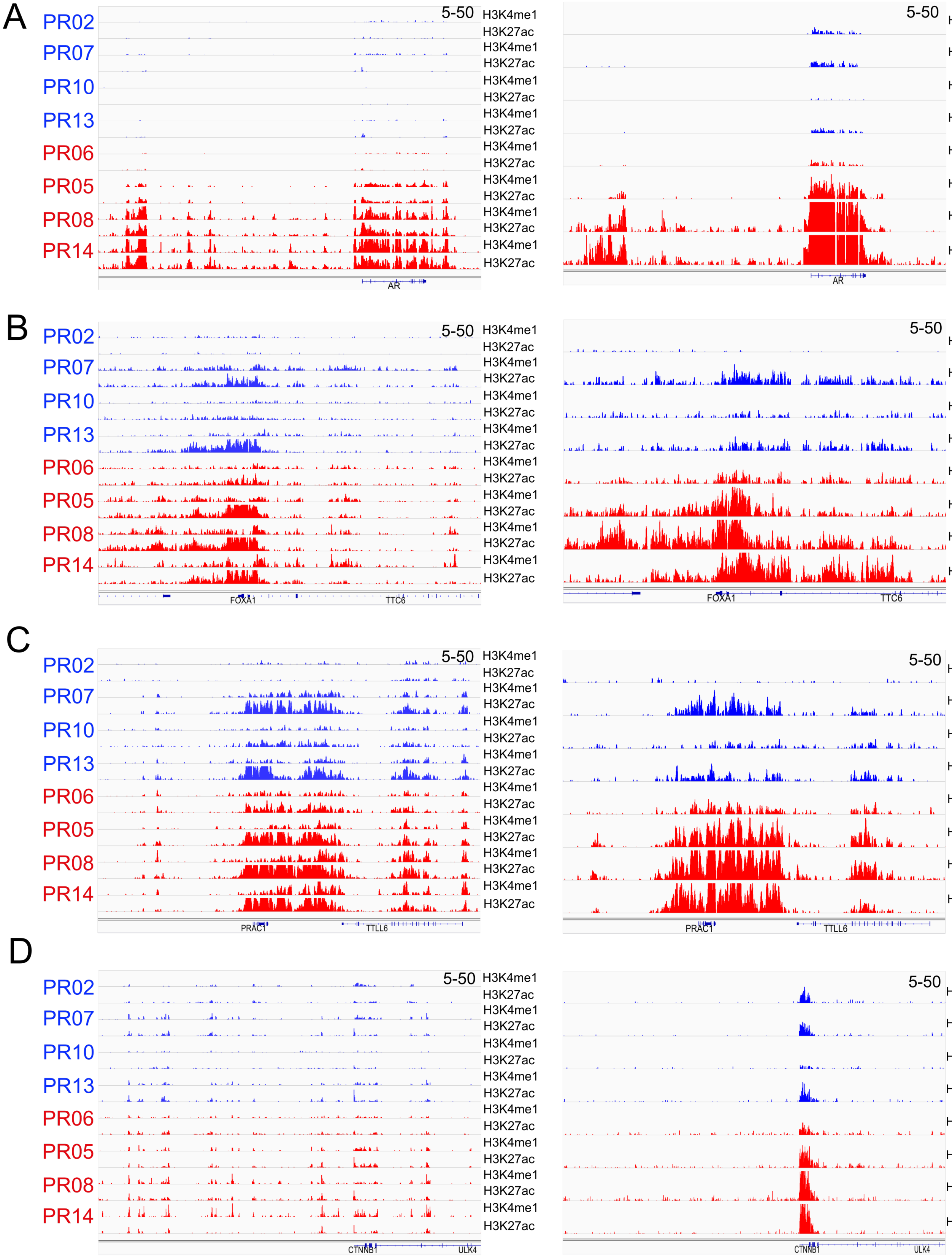


**Figure S5: Genome browser views of H3K4me1, H3K27ac and H3K79me2 in naive and therapy-resistant prostate tumors**

A-D) ChIP-seq tracks displaying individual H3K4me1, H3K27ac (left) and H3K79me2 (right) histone modifications on the AR, B) FOXA1, C) HOXB13 and D) CTNNB1 loci in naive and resistant tumors.

Figure S6

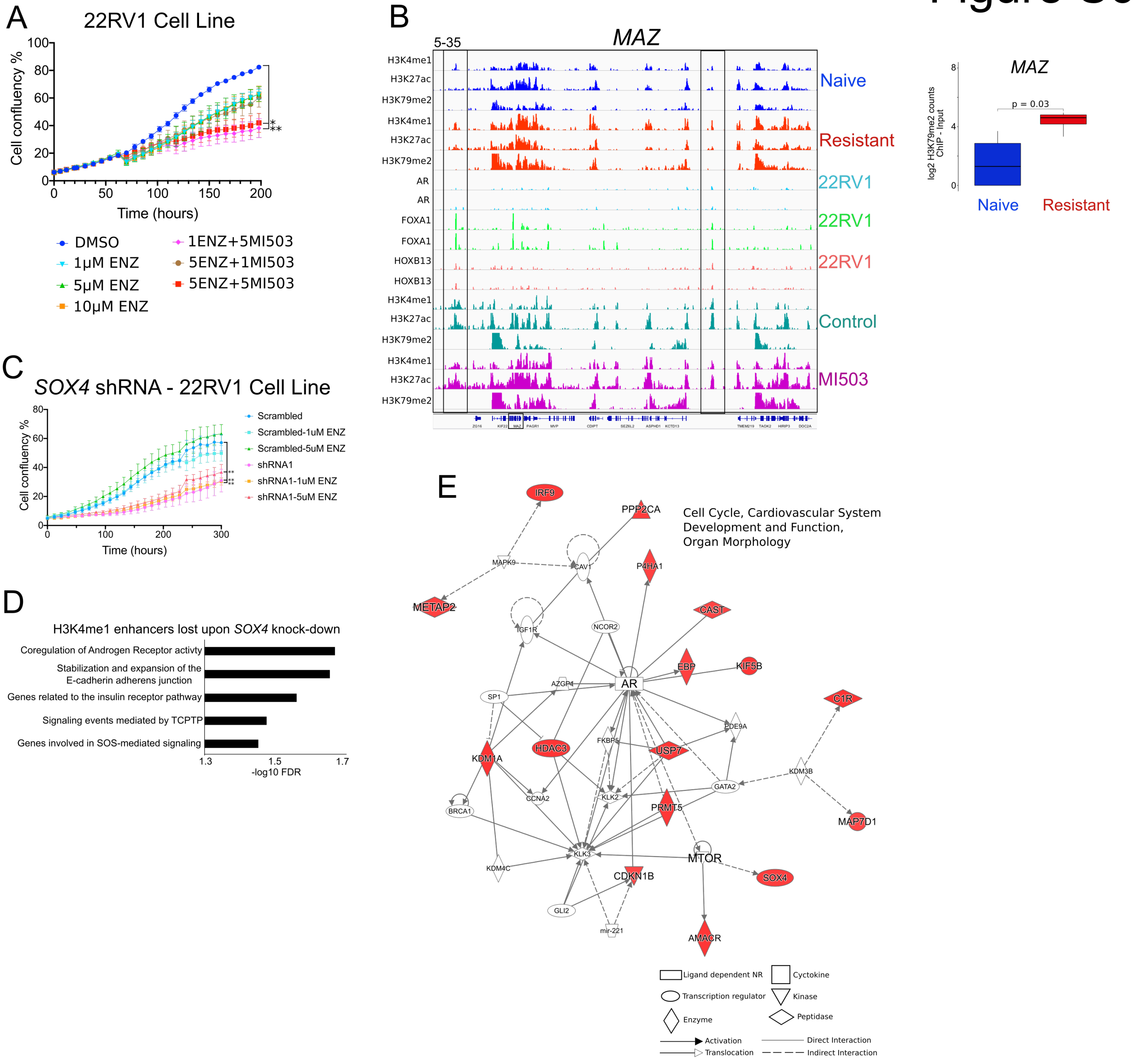


**Figure S6: SOX4 is co-expressed with proliferative-driver genes in therapy-resistant prostate cancers**

A) Representative proliferation assays in 22RV1 cell lines treated with DMSO (Control), enzalutamide (MV3100) or MV3100 + MI-503 inhibitor combinations for 48 hours. n = minimum of 3 biological and 3 technical replicates in each condition. P-values based on multiple comparison ANOVA between groups. * = $p < 0.05$, ** = $p < 0.005$

B) ChIP-seq tracks displaying H3K4me1, H3K27ac and H3K79me2 in naive and resistant tumors, the AR, FOXA1 and HOXB13 TF from WT 22RV1 cells (Kron et al., 2017), and H3K4me1, H3K27ac and H3K79me2 in WT 22RV1 cells treated with DMSO (Control) or 10μM MI-503 on the MAZ locus (left). Black bar denotes putative H3K4me1-associated developmental enhancers that are lost upon 10μM MI-503 treatment. ChIP-seq count data (log2) displaying H3K79me2 for TF identified in enhancers gained in resistant tumor samples that are also expressed within patient tumor samples treated with ABI-ENZ (>20 FPKM) from the SU2C-PCF dataset (Abida et al., 2019) (right).

C) Proliferation assays in Control (Scrambled) or SOX4 shRNA (shRNA-A3 and A4) 22RV1 cell lines treated with DMSO (Control) or MDV-3100 treatment. P-values represent pairwise t-test comparison between control and SOX4 shRNA in the presence or absence of MDV-3100. ** = $p < 0.005$. n = minimum of 2 biological and 4 technical replicates in each condition.

D) Top significant GREAT MSigDB Pathways based on Control (Scrambled) H3K4me1 enhancer regions (outside -/+5kbTSS) in 22RV1 cells. Hyper FDR q-values are displayed as -log10.

E) IPA gene network analysis based on the top -/+100 genes co-expressed with SOX4 in naive tumor samples from the SU2C-PCF dataset (Abida et al., 2019). Legend specifies potential gene relationships. Red indicates gene is directly expressed together with SOX4.
